# Phylogenomic timetree-calibrated speciation clocks for *Caenorhabditis* nematodes reveal slow but disproportionate accumulation of post-zygotic reproductive isolation

**DOI:** 10.1101/2025.06.18.660443

**Authors:** Daniel D. Fusca, Maia N. Dall’Acqua, Santiago Sánchez-Ramírez, Asher D. Cutter

## Abstract

Reproductive isolation and genomic divergence both accumulate over time in the formation and persistence of distinct biological species. The pace of “speciation clocks” quantified with pre-zygotic and post-zygotic reproductive isolation, however, differs among taxa, with pre-zygotic isolation tending to evolve sooner in some but not all taxa. To address this issue in nematodes for the first time, here we infer the species tree and divergence times across the phylogeny of 51 species of *Caenorhabditis*. We incorporate several molecular evolutionary strategies in phylogenomic dating to account for complications in this group due to lack of fossil calibration, deep molecular divergence with synonymous-site saturation, and codon usage bias. By integrating divergence times with experimental data on pre- and post-zygotic reproductive isolation, we infer that post-zygotic isolation accumulates faster than pre-zygotic isolation in *Caenorhabditis* and that hybrid sterility evolves sooner than hybrid inviability. These findings are consistent with speciation being driven principally by intrinsic isolating barriers and the disproportionate fragility of germline developmental programs to disruption. We estimate that it takes approximately 50 million generations for intrinsic post-zygotic reproductive compatibility to be reduced by half, on average, between diverging pairs of *Caenorhabditis*. The protracted reproductive isolation clocks in *Caenorhabditis* may, in part, reflect the capacity to retain population genetic hyperdiversity, the incomplete sampling of global biodiversity, and as-yet uncharacterized incipient or cryptic species.

## Introduction

The tempo of speciation is determined by the combined rate at which different reproductive isolation barriers accumulate between populations over time [1, 2]. Isolating barriers can be composed of both extrinsic environmental factors and intrinsic developmental genetic factors [3, 4]. In turn, these factors influence the propensity for mating to take place at all or for fertilization to occur in the presence of mating (pre-mating and post-mating pre-zygotic reproductive isolation, respectively, together comprising pre-zygotic isolation), and for the potential of hybrid individuals to survive and reproduce successfully (post-zygotic reproductive isolation) [5]. A central challenge in understanding the production of biodiversity through the speciation process is to resolve the relative importance, mechanisms of action, and pace of accumulation of these diverse components that cause reproductive isolation in the origin and maintenance of distinct species.

One difficulty to achieving consensus on these central questions arises due to disparities among taxa in the rates at which different reproductive isolation barriers accumulate: organisms often show distinct “speciation clocks” [6–10]. The disparities, however, are not simply quantitative differences in the rate per unit time. Taxa differ in whether extrinsic or intrinsic post-zygotic barriers tend to accumulate first and in whether post-zygotic or pre-zygotic barriers tend to accumulate first [5, 11, 12]. For example, post-zygotic isolation tends to evolve first in killifish, stalk-eyed flies, copepods, and bellflowers among other taxa [13–16] whereas, as more commonly reported, the evolution of pre-mating barriers precede post-zygotic barriers in *Drosophila*, centrarchid fish, and many other animals and plants [5, 6, 17, 18]. Such differences may implicate divergent ecological adaptation, reproductive character displacement, or developmental system drift as especially important in some systems relative to others. While post-zygotic reproductive isolation tends to first impact fertility (relative to viability) [5, 8, 19] and the development of earlier phases of the life cycle [18, 20, 21], consideration of broad trends of causal developmental mechanisms in the speciation process remain unresolved [22, 23]. Taxa also vary substantially in the absolute time, in generations or years, required to achieve complete reproductive isolation [2, 12, 24–26], despite some consistency in mean time to speciation across groups [27]. To build toward a consensus on the causes of such heterogeneity, it is instructive to characterize “reproductive isolation clocks” in diverse and neglected groups of organisms and across distinct barrier types.

*Caenorhabditis* nematode roundworms provide a powerful model system for experimental biology [28] and comparative evolutionary biology [29–32]. Consideration of the nematode phylum, however, is underrepresented in studies of speciation [24], in part due to relatively recent discovery and attention to species that demonstrate incomplete reproductive isolation to permit experiments involving hybrids [33–48]. Limited diagnostic morphological trait variation constrains the discrimination of species in these non-parasitic and bactivorous organisms [42, 49], making post-zygotic reproductive isolation itself an invaluable diagnostic criterion [36].

Consequently, studies of reproductive isolation among species of *Caenorhabditis*, to date, have focused primarily on intrinsic post-zygotic isolation, demonstrating evidence for Haldane’s rule in the form of stronger sterility and inviability among hybrid males, as well as cross-dependent asymmetries in hybrid dysfunction (Darwin’s corollary to Haldane’s rule, [50]) and intra-specific genetic variation for the magnitude of reproductive isolation [33, 51–53]. The mechanism of hybrid male sterility, in one instance, depends to a large degree on misregulation of small RNAs and spermatogenesis due to multi-locus X-linked factors [35, 47, 54, 55]. Hybrids show profound sex differences in gene misregulation [56], and hybrid inviability manifests disproportionately prior to hatching [20, 39, 45], in part due to differential expansion of F-box genes in one pair of species [44]. Moreover, post-mating pre-zygotic barriers are evident between some species as a result of sperm migration defects that can also cause reproductive interference in multi-species communities, though assessments of pre-mating isolation are largely uncharacterized [57–60].

Given sampling within the *Caenorhabditis* genus that now exceeds four score species [61], combined with substantial genome resources for them [29, 62] and standardized approaches to assessing biological species identity [36], this group is positioned to contribute substantively to core problems of speciation. The great span of genomic sequence divergence across *Caenorhabditis* species and lack of fossil record, however, present a challenge to dating the history of speciation in the genus to place molecular evolution, trait change, and reproductive isolation in a robust temporal framework [63]. With these issues in mind, here we integrate several strategies to date the times to most recent common ancestors across a phylogeny of 51 species of *Caenorhabditis* to provide estimates of speciation times across the genus. We then relate the time since speciation on the resulting timetree to metrics of reproductive isolation for over 250 species pairings to document the evolutionary pace of intrinsic pre-zygotic and post-zygotic reproductive isolation for *Caenorhabditis* nematodes.

## Results

### Divergence time estimation creates a *Caenorhabditis* timetree

We first constructed a phylogenomic species tree for 51 species of *Caenorhabditis* nematode roundworms from 3130 single-copy gene trees (Figure 1). This tree topology, with minor alterations, was robust to methodological effects of including only highest quality ortholog alignments (n=1768 gene trees), concatenation of orthologs (n=1 gene tree), inclusion of multicopy orthogroups (n=9926 gene trees), and simultaneous inference of tree topology and divergence times (n=51 gene trees). The species tree also supports the key findings of recent phylogenies generated for subsets of these species, including the composition of major clades and nodes exhibiting challenges with concordance [29, 62, 64, 65]. Most nodes in the phylogeny (45/50 = 90%) were confidently resolved, with just 5 internal nodes contributing to alternative topologies (Figure 1). In particular, 7 of the 50 internal nodes showed low site concordance factors < 33.3%, indicative of substantial gene tree – species tree discordance potentially resulting from incomplete lineage sorting in these regions of the phylogeny, with a subset of 3 of these nodes also showing a posterior probability <1 (Figure 1) (Supplementary Figure S4).

**Figure 1.**
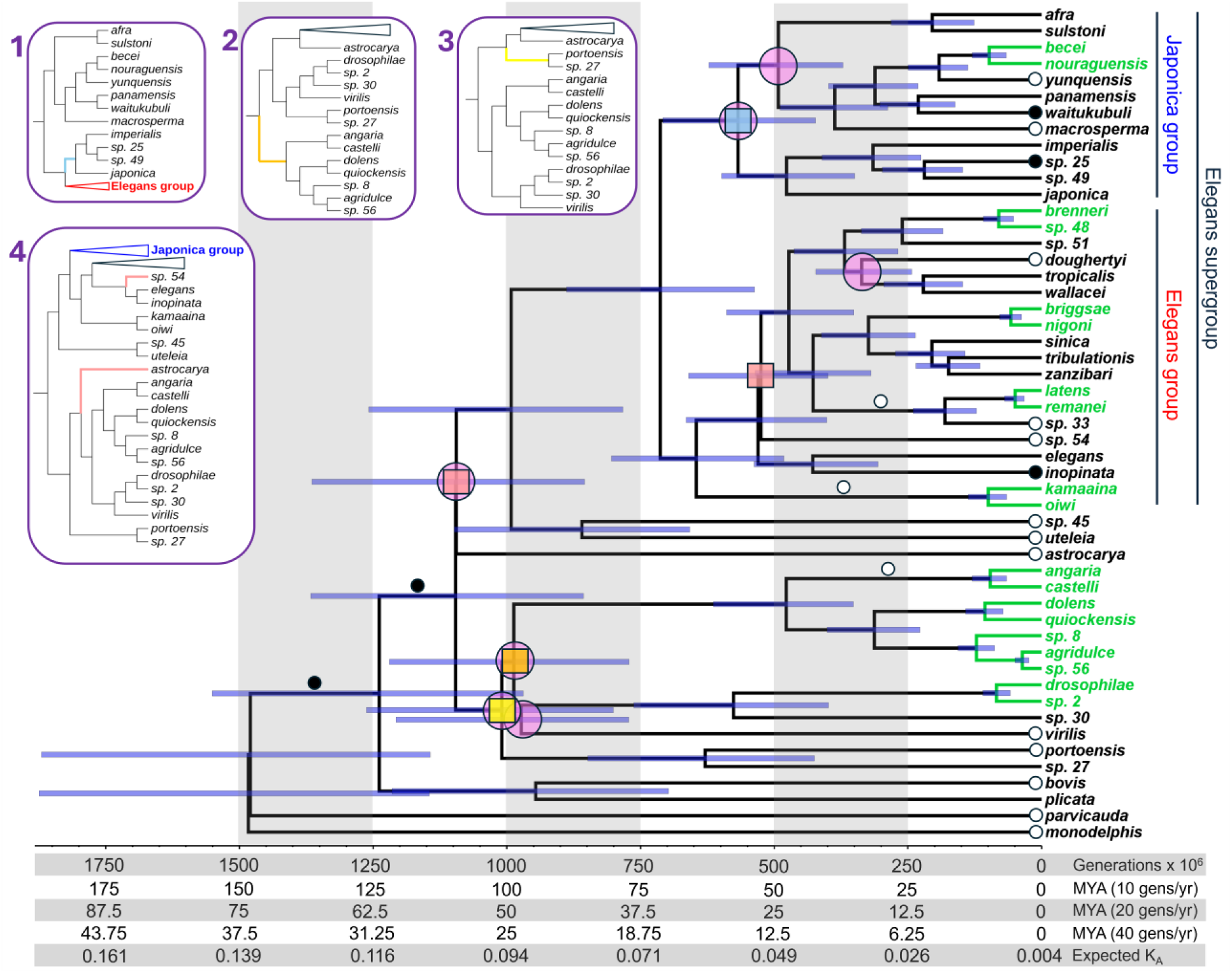
Dated phylogeny of the *Caenorhabditis* genus. Divergence times are provided in units of millions of generations (assuming a constant generation time across the phylogeny), years (assuming generation times of 10, 20, or 40 generations per year for the calibration species), and non-synonymous site substitutions (lineage-specific *K*_A_, i.e. pairwise *K*_A_/2 between a given species pair; derived from our regression between divergence time and median pairwise *K*_A_ value). Clades in green indicate the 10 speciation events used as calibration points for dating. Error bars (blue) indicate the 95% Highest Posterior Density intervals for each date estimate. Insets show the 4 alternate tree topologies, highlighting with distinct colors the branches with alternate topologies; corresponding colored squares on the primary topology indicate the nodes contributing to these 4 alternate topologies. Pink circles indicate nodes with site concordance factors < 33.3%. Black and white circles indicate branches with significantly elevated (black) or reduced (white) relative substitution rates (see Supplementary Figure S4).

We next sought to use the *Caenorhabditis* species tree to infer times of common ancestry among the species. The lack of fossil record for *Caenorhabditis*, however, makes this goal challenging, especially when coupled with saturated synonymous-site divergence for >96% of species contrasts among the 51 species in our analysis (Supplementary Figure S1, Supplementary Table S3). Consequently, we aimed to calibrate divergence times by integrating a mutation accumulation estimate of mutational input [66] with putatively neutral divergence between the most-closely related sets of species. Specifically, we identified 10 nodes on the species tree to use as calibration points (synonymous-site substitution rate *K*_S_ < 0.4). To obtain the best possible estimates of neutral divergence between them [63], we accounted for multiple mutational hits for large numbers of 1:1 ortholog pairs per species (n=8774 to 15,204 orthologs) and further corrected sequence divergence of synonymous sites of orthologous genes (*K*_S_) for its correlation with codon usage bias due to weak but influential selection on synonymous sites (effective number of codons, ENC; *P* < 2.2 x 10^-16^ for all species pairs, Spearman’s rank correlation) (Figure 2A, B, Supplementary Table S4). We then converted these distributions of corrected *K*_S_ values (*K*_S_′) to distributions of divergence times under a strict molecular clock for use as calibration time priors (Figure 2C) (Supplementary Figure S2), using an experimentally determined nucleotide mutation rate estimate from *C. elegans* [66]. Using these 10 calibration node time distributions as priors (Supplementary Figure S2), the species tree topology, and DNA coding sequence alignments for 205 single-copy ortholog groups, we performed Bayesian phylogenomic dating in BEAST to estimate divergence times across the *Caenorhabditis* phylogeny in a manner that made use of amino acid sequence divergence.

**Figure 2.**
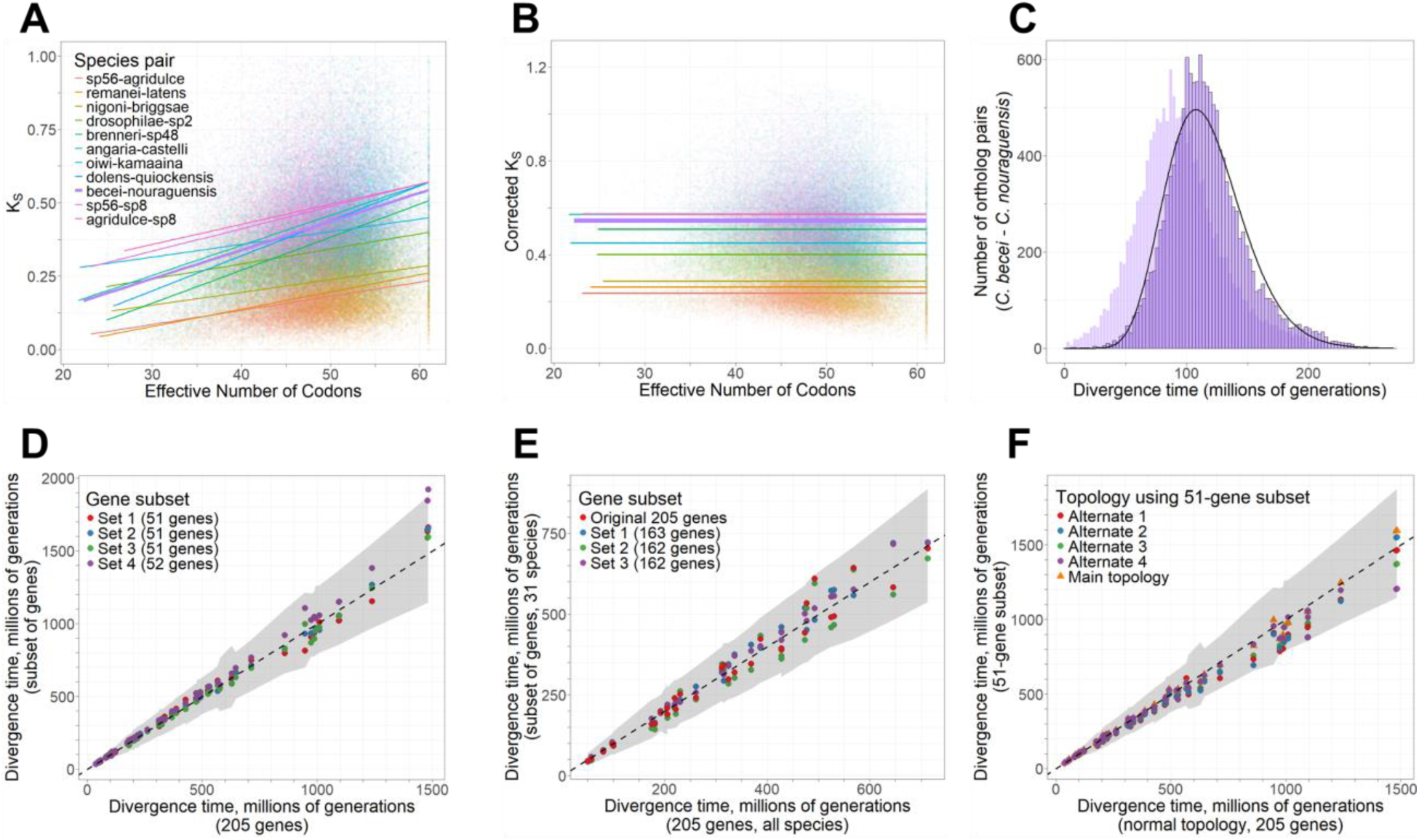
Divergence time calibration and tests of sensitivity of date estimates. **A.** Positive correlation between Effective Number of Codons and uncorrected synonymous-site substitution rate *K*_S_ for 1:1 orthologs of 11 calibration species pairs. **B.** No relationship between Effective Number of Codons and corrected *K*_S_ (i.e. *K*_S_′) for 1:1 orthologs of 11 calibration species pairs. Lines in panels A and B indicate least-squares regression lines for each species pair; thick purple line indicates *C. becei* – *C. nouraguensis* shown in panel C. **C.** Distribution of divergence times for the example species pair of *C. becei* and *C. nouraguensis*, based on ENC-corrected values of *K*_S_′ (foreground, black-outlined bars) or uncorrected values of *K*_S_ (background, lighter bars) for 1:1 orthologs to divergence times using the strict molecular clock application of mutation rate estimate. Black curve shows the density of the prior distribution from fit to a gamma distribution that was input to BEAST for this calibration point. Supplementary Figure S2 shows distributions and density fits for all species pairs used in calibration. **D.** Comparison of divergence time estimates for each species pair between the primary dates based on 205 genes and the 4 subsets of these genes. Shaded interval in D-F gives the 95% Highest Posterior Density (HPD) interval around the 205-gene dates. Note that while one point from Set 4 falls outside the shaded interval; the 95% HPD interval on this subset date (not shown) overlaps with the shaded interval. **E.** Comparison of divergence time estimates for each species pair in the Elegans supergroup using either the primary dates based on 205 genes (considering all 51 species) or the 4 subsets of the 692 Elegans supergroup genes (considering only the 31 Elegans supergroup species). **F.** Divergence time estimates for each species pair contrasted for estimates from the 205 genes and the main species tree topology versus dates based on a subset of 51 genes using 5 different tree topologies (primary topology and 4 alternates indicated in Figure 1). Dashed lines in panels D-F gives the 1:1 line.

By applying this novel approach, we estimate that the 51 *Caenorhabditis* species considered here shared a common ancestor approximately 1.48 billion generations ago (95% Highest Posterior Density interval: 1.15 – 1.88 billion generations ago) (Figure 1, Supplementary Table S3). For an assumption of 20 effective generations per year (Xu Wei and Marie-Anne Felix, personal communication), this timing would correspond to a 74 Mya Late Cretaceous last common ancestor for *Caenorhabditis* (Figure 1). While an assumption of faster effective generation time per year will lead to more recent divergence date estimates in units of years (Figure 1), it will not influence the date estimates scaled to time in units of generations. We also infer that the set of 31 species in the Elegans supergroup, which includes *C. elegans*, has diversified as a distinct clade within the genus for about half of the evolutionary duration of the genus as a whole, since approximately 713 million generations ago. Lineage-through-time (LTT) plot analysis shows that the accumulation of *Caenorhabditis* lineages over time does not deviate significantly from the expectations of log-linear growth (Pybus & Harvey’s γ = −1.1749, *P* = 0.24) (Figure 3A). We observe a nominal excess in lineage accumulation approximately 1 billion generations ago, however, coinciding with several rapid (but highly discordant, in terms of the underlying sequence alignments) speciation events in a clade of early-diverging species (Figure 3B, Supplementary Figure S5). Comparison of the observed accumulation of lineages through time to the average expected under a growth model with known undersampling of species diversity reveals a slight plateau in the LTT curve toward the present (Figure 3B), suggesting that species unsampled in our phylogeny will disproportionately contribute to short branch lengths and closely-related species pairs.

**Figure 3.**
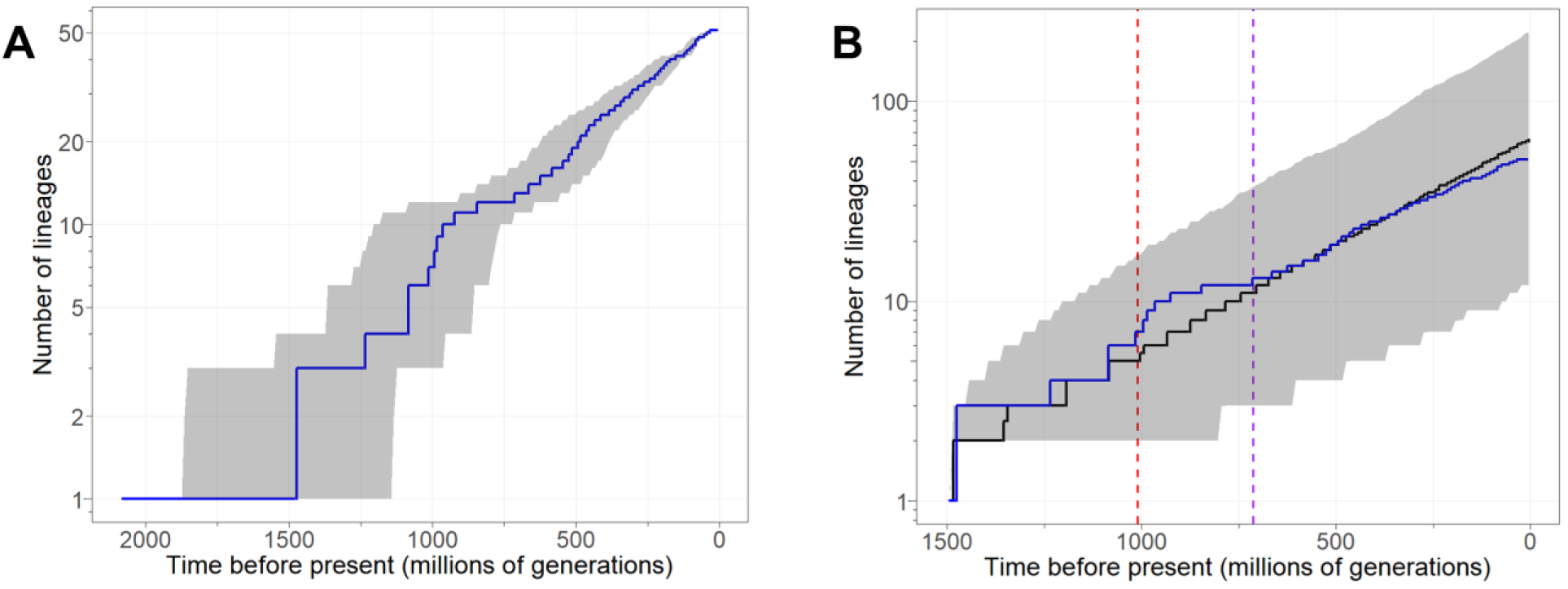
Lineage-through-time plot for the *Caenorhabditis* genus. **A.** Observed cumulative number of lineages through time in blue for the 51-species phylogeny. Shaded intervals give the 95% upper and lower quantiles around the median, based on the posterior distributions of divergence times calculated by BEAST. **B.** Comparison of observed (blue) and expected (black) lineage accumulation curves. Expected lineage accumulation from 1000 random phylogenies with the same total age as our primary phylogeny, but any final number of tips. Phylogenies were generated under the birth rate (λ=0.0248 per lineage per 10 million generations) estimated from our primary phylogeny when assuming a species sampling fraction of 59% (51/86). Shaded intervals give the 95% upper and lower quantiles around the median, based on the 1000 random trees. Vertical lines indicate the ages of the Elegans supergroup (purple) and the clade defined by the most recent common ancestor of *C. angaria* and *C. portoensis* (red).

Our dating analysis also indicates that *C. elegans* diverged from its nearest known relative (*C. inopinata*) approximately 428 million generations ago, and approximately 530 million generations ago from *C. briggsae* (Figure 1) (Supplementary Table S3). We estimate the most recent divergence time between known species of *Caenorhabditis* to be 36 million generations ago (*C. agridulce* and *C*. sp. 56). Deeper nodes in the phylogeny are associated with broader confidence intervals on date estimates. We also note that these divergence date estimates do not account for ancestral polymorphism, and so represent slight overestimates of species split times [67]. Although neutral polymorphism is unknown for most extant species – and all extinct ancestral species – the typically high polymorphism (θ) observed in outbreeding *Caenorhabditis* [68, 69] suggests that split times may commonly be ∼10 million generations (i.e., ½θ_neutral_/µ generations) more recent than the BEAST estimates imply.

To test the sensitivity of dating to gene composition, we repeated our dating procedure on four subsets of ∼50 genes compared to the full 205 gene set. With the exception of one node in one gene subset, all node date estimates fell within the 95% Highest Posterior Density interval for the 205-gene set (Figure 2D), consistent with our dating procedure being robust to ortholog composition. Similarly, after repeating our dating procedure for an expanded set of 692 groups of single-copy orthologs restricted to the 31 species in the Elegans supergroup clade, all divergence time estimates again fell within the 95% Highest Posterior Density intervals of the 51-species dates estimated from the 205-gene set (Figure 2E). We also tested the sensitivity of dating to four alternate possible topologies of the species tree (Figure 1), based on a subset of 51 single-copy genes. The divergence times estimated using these alternate topologies always fell within the 95% Highest Posterior Density interval of our primary divergence time estimates (Figure 2F), consistent with divergence time estimates being robust to possible misspecifications of the underlying tree topology.

Tests for rate heterogeneity across the tree identify 16 of the 51 terminal branches to exhibit modest differences from the average rate of molecular evolution across the phylogeny (Figure 1) (Supplementary Figure S4). Of these 16 cases, 3 showed rates with 95% Highest Posterior Density intervals exceeding the global average and 13 showed rates lower than the global average. In addition, 5 internal branches showed rates of molecular evolution that differed from the global average, with 2 such nodes near the base of the phylogeny exhibiting elevated rates (Supplementary Figure S4). Such rate heterogeneity might arise from a number of possible biological sources, including mutation rate differences, effective generation time differences among lineages, or differences in effective population size that affects the fraction of neutral substitutions in amino acid divergence.

### A simple method for *Caenorhabditis* divergence time estimation

As new *Caenorhabditis* species continue to be discovered and sequenced at a rapid pace [29, 36, 61], we wondered whether it was possible to use our estimated divergence times to predict divergence times for new species pairs. We found that simple calculations of median *K*_A_ from orthologs between species increase linearly with the BEAST-derived divergence time estimates (linear regression adjusted R^2^ = 0.983, *P* < 2.2 x 10^-16^) (Figure 4A, Supplementary Table S3), which are more onerous to calculate. This finding supports the idea that divergence times for new species can be well-approximated from *K*_A_ values, which can be calculated much more quickly and easily than with Bayesian dating. To explore this idea concretely, we obtained genome sequences for *C. auriculariae* and *C. niphades* (neither of which were included in our original set of 51 species) and calculated median *K*_A_ between these species and the other 51 species using 1:1 orthologs (Supplementary Table S5). Consistent with the placement of *C. auriculariae* as sister to *C. monodelphis* near the base of the genus, application of our *K*_A_– divergence time regression implies that *C. auriculariae* diverged from *C. monodelphis* about 1.07 billion generations ago, and from the rest of the genus approximately 1.38 – 1.52 billion generations ago (Figure 4B, Supplementary Table S5). These latter dates are fully consistent with our Bayesian estimated age of 1.48 billion generations for the root of the genus. Our phylogenetic placement of *C. niphades* and calculations of *K*_A_ imply divergence from its sister clade sometime 521 – 576 million generations ago (Figure 4C; Supplementary Figure S3, Supplementary Table S5), consistent with our Bayesian divergence time estimates for the Japonica Group. Bayesian and *K*_A_-derived estimates with *C. niphades* also are consistent for the root of the genus and the origin of the Elegans supergroup (Figure 4C). Overall, we find that a simple relationship between non-synonymous sequence divergence and Bayesian divergence time estimates for our 51 species can be applied to infer reasonable divergence times for other *Caenorhabditis* species simply by substituting in median pairwise *K*_A_ values to the equation *T* = [(*K*_A_ – 7.888×10^−3^) / (1.798×10^−10^)] / (µ* / 2.33×10^−9^) generations, where µ* corresponds to the neutral mutation rate assumption (which may differ from ours). Divergence time estimation from average *K*_S_ values, after correction for multiple hits and codon usage bias, also ought to be appropriate for species pairs with *K*_S_′ < 1 (i.e. *T* < 250 million generations; Figure 1, Supplementary Figure S1B).

**Figure 4.**
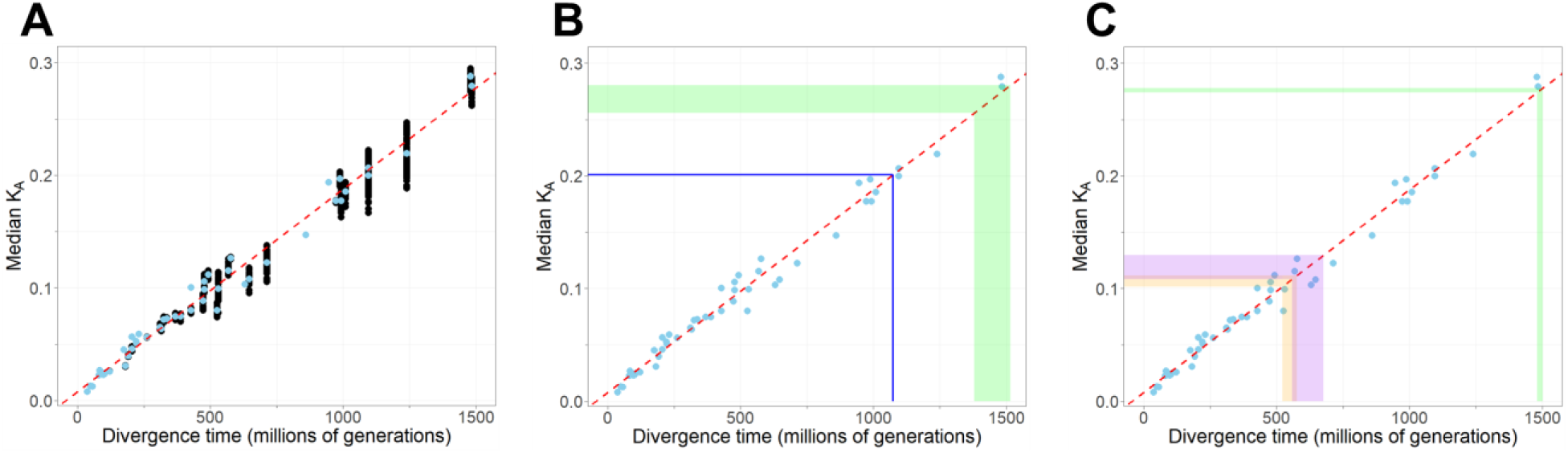
Non-synonymous site divergence (*K*_A_) relates positively with estimates of divergence time. **A.** Comparisons based on 1:1 orthologs, with median *K*_A_ across orthologs for a given species pair indicated by black points for all unique pairs of the 51 species. For each node on the species tree, blue circles indicate the median value of median *K*_A_ taken across all species pairs at that node (including nodes with just a single species pair) in panels A-C. Least-squares regression line fit (red) to the 50 per-node median values in panels A-C. **B.** Predicted divergence times involving *C. auriculariae* shown with colored lines intersecting the x-axis indicate the upper and lower bounds of predicted divergence times, derived from the corresponding *K*_A_ ranges of the same color. Blue line indicates the *K*_A_ between *C. auriculariae* and *C. monodelphis*, and green band indicates the range of *K*_A_ between *C. auriculariae* and every species besides *C. monodelphis*. **C.** Predicted divergence times involving *C. niphades*. Orange band indicates the range of *K*_A_ between *C. niphades* and species in its sister clade, purple band indicates the range of *K*_A_ between *C. niphades* and species in the Elegans group, and green band indicates the range of *K*_A_ between *C. niphades* and the earliest-diverging species (*C. monodelphis* and *C. auriculariae*).

### Extensive post-zygotic reproductive isolating barriers despite incomplete pre-zygotic isolation between most *Caenorhabditis* species

Because fitness-related reproductive barriers provide a key diagnostic criterion in delimiting species of *Caenorhabditis* that often are morphologically cryptic [36, 49], inter-species crossing experiments provide a rich source of information about the magnitude of reproductive isolation in this group. We computed metrics of reproductive isolation for 258 published inter-species pairings (19% of the 1332 possible reciprocal pairwise combinations of the 37 *Caenorhabditis* species represented) and 5 within-species pairings to assess the degree of pre-zygotic reproductive isolation due to mating or gametic barriers, post-zygotic reproductive isolation in F1 hybrids in terms of viability and developmental progression, as well as F2 hybrid breakdown (Supplementary Table S2). By integrating these indices of the magnitude of reproductive isolation with the evolutionary distance between species pairs, we then document the prevailing temporal trends in the accumulation of isolating barriers in species formation for *Caenorhabditis*.

Despite the vast majority of *Caenorhabditis* inter-species crosses showing incomplete pre-mating isolation (96.3% of pairs showed mating between species among 136 pairings that assessed mating; 53.9% of 243 crosses, including unknown mating status), the vast majority (98.4%, 239 of 243 pairs) also show complete reproductive isolation in terms of failure to fertilize or post-zygotic hybrid inviability or sterility (Figure 5). Studies to date, however, may underestimate the magnitude of pre-mating isolation because (1) they are typically performed in no-choice mating arenas without the option to discriminate between conspecific and heterospecific mates and (2) 37.0% of reports of crossing experiments are ambiguous as to whether the terminal isolating barrier is failure to mate, failure to fertilize, or embryonic arrest. Fertilization was reported to not occur in nearly one quarter (22.6%) of inter-species crosses, however, indicative of frequent and substantial pre-mating and/or post-mating pre-zygotic isolating barriers. Two documented mechanisms of post-mating pre-zygotic isolation include failure to inseminate upon mating [70] and ectopically-invasive sperm that induce heterospecific female sterility and mortality [57].

**Figure 5.**
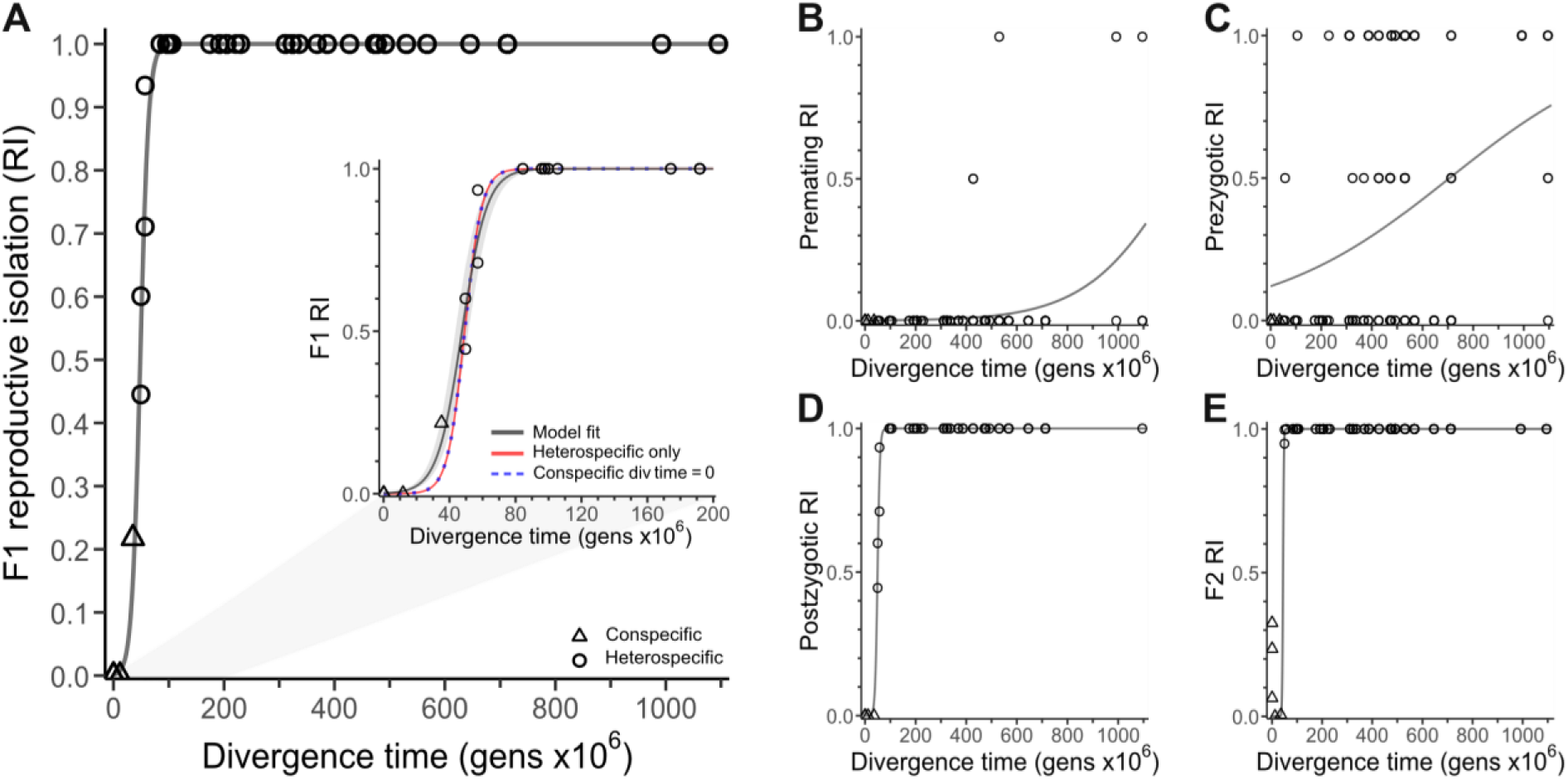
Reproductive isolation (RI) clocks for *Caenorhabditis*. **(A)** Logistic function fit to overall F1 RI between all species pairs, with coalescent time depths used for within-species data points. Inset shows logistic fits with conspecific divergence times set to 0 (dashed blue line) or excluded (red line). RI values of 1 indicate that no fertile F1 offspring are produced. **(B)** Logistic function fit to pre-mating RI as a function of divergence time. Values of 1 indicate that mating is not observed. **(C)** Logistic fit to pre-zygotic RI as a function of divergence time. Values of 1 indicate complete pre-mating or post-mating pre-zygotic isolation that prevents the production of embryos; RI values of 0.5 indicate documented cases of gametic isolation. **(D)** Logistic fit to post-zygotic RI as a function of divergence time. Values of 1 indicate that all F1 hybrids are sterile or inviable. **(E)** Logistic fit to F2 RI as a function of divergence time. Values of 1 indicate that F2 offspring are not produced or are all sterile or inviable. Logistic function fits for each component of RI were fit using all species-pair data; subsample with phylogenetically independent contrasts in Supplementary Figure S7.

Embryonic arrest represents the terminal hybrid phenotype in at least 33.3% of cases, with only 7% of inter-species crosses reporting hybrid development to larval or adult stages (Figure 6). Gastrulation appears especially sensitive to embryonic arrest, though its precursors are detectable at the one-cell stage in some cases [45] and inter-species hybrids differ in the distribution of embryonic stages experiencing arrest [39, 51]. In one species pair that yields adult hybrids, larvae experience little inviability despite substantial developmental arrest in embryos [20]. In only two species pairs are fertile adult F1 hybrids characterized: *C. remanei - C. latens* (both hybrid sexes fertile with maternal *C. latens*, females fertile with maternal *C. remanei*) and *C. briggsae - C. nigoni* (only female hybrids fertile in both cross directions) [33, 34, 37] (see also new cases in [61]).

**Figure 6.**
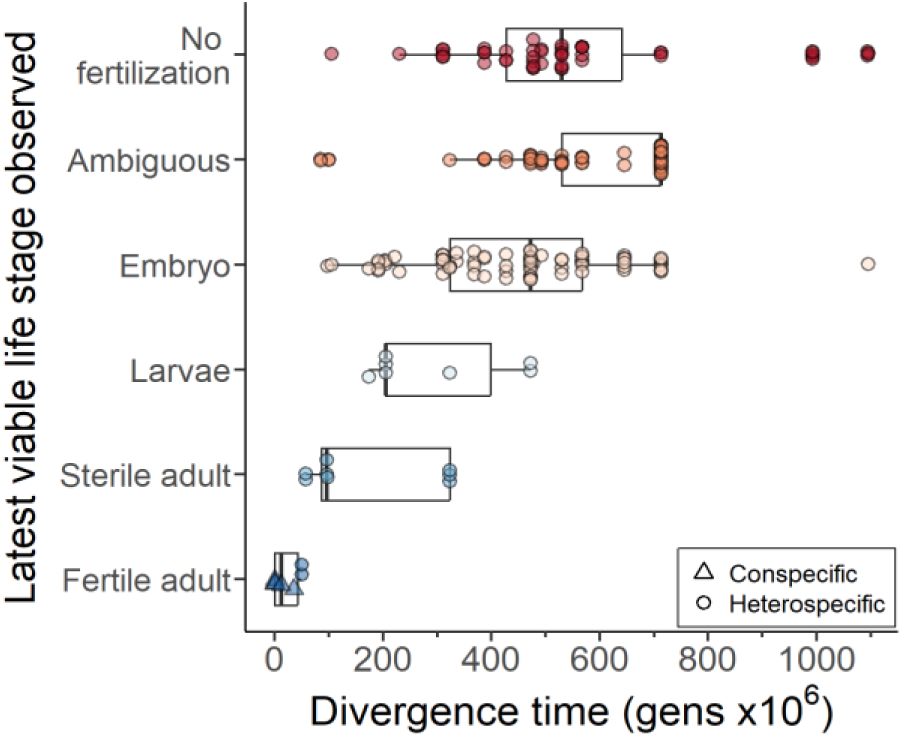
Terminal stage of development achieved in F1 hybrids. Latest observed viable stage of F1 hybrids reported in the literature, with reciprocal cross-directions each plotted when available; “ambiguous” denotes cross reports that did not indicate whether eggs were fertilized or unfertilized. Latest viable life stage observed is positively correlated with divergence time (Spearman’s ρ = 0.43, P < 1×10^−11^). Subsample analysis using phylogenetically independent contrasts and averages of reciprocal crosses in Supplementary Figure S8.

### Reproductive isolation clocks record faster evolution of post-zygotic than pre-zygotic isolation in *Caenorhabditis* speciation

We next integrated the metrics of reproductive isolation between species pairs with the age of the phylogenetic node of their most recent common ancestor. By fitting logistic functions, we characterized the stereotypical profile of accumulation of reproductive isolation to estimate the divergence time required to yield 50% reproductive isolation for each metric (Figure 5; Table 1). We estimate that 47.6 million generations accrue on average for 50% overall reproductive isolation to manifest at F1 and 43.8 million generations in F2. Inclusion of within-species inter-population measures did not appreciably affect these estimates (Supplementary Table S2). We also found that post-zygotic isolating barriers accumulate >70-fold faster than pre-zygotic barriers in general and >30-fold faster than pre-mating barriers to reproductive isolation (*b*_post-zyg_ / *b*_pre-zyg_ or *b*_post-zyg_ / *b*_pre-mating_; Figure 5), yielding estimated times to 50% pre-zygotic isolation requiring over 700 million generations (Table 1). Partitioning F1 post-zygotic isolation by the latest developmental stage achieved by hybrids shows a positive correlation between divergence time and the magnitude of developmental disruption (phylogeny-uncorrected Spearman’s ρ = 0.43, P < 1×10^−11^), such that the terminal developmental stage achieved by F1 hybrids occurs earlier in ontogenetic development for those species pairs separated by greater spans of evolutionary time (Figure 6). In particular, the average divergence time between species pairs that exhibit embryonic hybrid inviability is longer ago than for those that show hybrid sterility (Wilcoxon rank sum test, P<0.01; Figure 6), consistent with the more rapid evolution of hybrid sterility than inviability.

**Table 1.** Logistic function fits of reproductive isolation (RI) over time.

| RI class | All comparisons |  |  |  | Phylogenetic independent contrasts |  |  |  |
| --- | --- | --- | --- | --- | --- | --- | --- | --- |
| | a | b | $T_{RI=0.5}^*$ | $T_{RI=0.95}^*$ | a | b | $T_{RI=0.5}$ | $T_{RI=0.95}$ |
| Premating | $9.62 \times 10^2$<br>$\pm 1.4 \times 10^3$ | $5.6 \times 10^{-3}$<br>$\pm 1.45 \times 10^{-3}$ | 1227.1 | 1753.1 | --- | --- | --- | --- |
| Prezygotic** | 7.3<br>$\pm 3.04$ | 0.003<br>$\pm 7.5 \times 10^{-4}$ | 707.5 | 1755.3 | 31.5<br>$\pm 65.74$ | $7.6 \times 10^{-3}$<br>$\pm 6 \times 10^{-3}$ | 454.3 | 842 |
| Postzygotic | $3.9 \times 10^4$<br>$\pm 2.77 \times 10^4$ | 0.214<br>$\pm 1.39 \times 10^{-2}$ | 49.4 | 63.2 | $1.4 \times 10^4$<br>$\pm 34.62$ | 0.19<br>$\pm 4.8 \times 10^{-5}$ | 49.2 | 64.3 |
| F1 | 595.6<br>$\pm 161$ | 0.134<br>$\pm 0.005$ | 47.6 | 69.1 | $1.41 \times 10^4$<br>$\pm 222.7$ | 0.19<br>$\pm 3.1 \times 10^{-4}$ | 49.1 | 64.3 |
| F2** | $4.15 \times 10^{11}$<br>$\pm 7.14 \times 10^{12}$ | 0.611<br>$\pm 0.35$ | 43.8 | 48.6 | --- | --- | --- | --- |

\* $T$ divergence time in gens $\times 10^6$ ; within-species divergence set to estimated coalescent time depths
\*\*Model fit failed to converge for phylogenetic independent contrasts of premating isolation and F2 reproductive isolation

These findings were robust to analysis of a subset of species pairs that were phylogenetically independent from other pairs. When considering only phylogenetically independent species pairs, the number of generations required to achieve 50% and 95% postzygotic and F1 reproductive isolation is comparable to the estimates that include all species pairs (Table 1, Supplemental Figure S7). However, pre-zygotic isolation is inferred to accumulate considerably faster when using the phylogenetically independent subsample (36% faster for *T*_RI_PreZ=0.5_ and 52% faster for *T*_RI_PreZ=0.95_) (Table 1). Divergence time and latest developmental stage retain a significant positive correlation when restricted to the subset of 16 species pairs that are phylogenetically independent (Spearman’s ρ = 0.65, P<0.01; Supplemental Figure S8).

## Discussion

### A *Caenorhabditis* “timetree”

Central to addressing many questions in biology is knowing the timescale over which evolutionary divergence accumulates between species and the pace at which biodiversity arises [27]. Despite the important role of *C. elegans* as a biological model organism [28] and the accumulating comparative resources from new species discoveries [29, 31, 36, 62], these timescale gaps in understanding have persisted for *Caenorhabditis* nematodes. In this study, we fill these gaps by generating estimates for dates of common ancestry for 51 *Caenorhabditis* species using a novel approach, by characterizing the pace of phylogenetic diversification, and by quantifying rates of accumulation of distinct components of reproductive isolation for the first time in the nematode phylum.

We generated a timetree for the *Caenorhabditis* phylogeny, including divergence time estimates for 51 species and their common ancestors. Our novel approach circumvents the major challenge to estimating dates of common ancestry shared by many soft-bodied organisms that lack fossils for temporal calibration [71, 72]. Here, we exploited direct experimental estimates of mutational input [66] to calibrate neutral substitution rates and divergence times between closely-related species. To generate the most accurate estimates for these calibration points, we accounted for molecular evolutionary complications due to synonymous-site mutational saturation between orthologs of most species pairs, as well as selection on synonymous sites from codon usage bias among highly-expressed genes. We then used these date estimate distributions as calibration points in Bayesian inference of dates deeper in the phylogeny from amino acid sequence divergence of shared orthologs. We are most confident in the dates measured in units of generations, as the ecologies of *Caenorhabditis* still have yet to reveal exactly how many generations a given species passes through each year in nature on average. Nonetheless, we propose that the genus as a whole may have originated approximately 74 Mya and that the closest known species pairs diverged within the last 1.8 million years.

Given that we anticipate ongoing discovery of many new species of *Caenorhabditis* alongside sequencing of their genomes, we sought to facilitate estimation of divergence times as new species become integrated into the phylogeny. Therefore, we derived a simple expression to relate our estimates of divergence time *T* derived from BEAST to the median values of *K*_A_ between gene orthologs (*T* = [(*K*_A_ – 7.888×10^−3^) / (1.798×10^−10^)] / (µ* / 2.33×10^−9^)). Future studies of additional species may thus calculate reasonable estimates of *T* for a given new ancestral node in a simple way without needing to conduct a full Bayesian analysis of the entire phylogeny.

### Why might completion of speciation be slow in *Caenorhabditis*?

We estimate the most recent dates of common ancestry for known *Caenorhabditis* species pairs at 36-50 million generations ago (median *K*_S_′ of 0.22 to 0.24, median *K*_A_ of 0.008 to 0.013) (Supplementary Figure S6), a deeper evolutionary timeframe than in many other animal taxa [73, 74]. This observation may reflect a truly protracted time to completion of speciation in this group, as suggested for some other organisms like amphibians [21, 74, 75], but also could represent a byproduct of the existence of unrecognized cryptic species or incomplete taxon sampling. It is plausible that completion of speciation might indeed proceed more slowly in this group, on average, given the enormous effective population sizes estimated for some *Caenorhabditis* species that yield exceptionally deep allele coalescent depths [76, 77]. For example, developmental system drift (DSD) can lead to the evolution of intrinsic reproductive isolation between allopatric populations on a timescale of *N_e_* generations [78]. *Caenorhabditis* species demonstrate widespread evidence for DSD [32, 56, 79–83]. Should DSD-mediated accumulation of post-zygotic genetic incompatibilities commonly drive speciation in this group, then the enormous *N_e_* exhibited by many *Caenorhabditis* species might provide one basis to why these organisms seem to show a slow rate of diversification compared to, e.g., mammals and birds. A scenario of *Caenorhabditis* typically undergoing speciation of allopatric populations also might explain the relatively weak pre-zygotic isolation despite strong post-zygotic isolating barriers between species [11]. A deeper understanding of the evolutionary processes underlying DSD over ontogeny may reveal how intrinsic post-zygotic reproductive isolation itself evolves [22, 23].

The morphologically-cryptic nature of different species of *Caenorhabditis* [34, 36, 49], however, also suggests the possibility that biologically distinct species might reveal themselves upon more detailed investigations of phenotypic reproductive isolation among wild isolate strains. Moreover, the subtle plateau of the lineage-through-time plot at the most recent timescales (Figure 3, Supplementary Figure S5) is consistent with the idea of incomplete taxon sampling [84], and new species discovery shows little sign of abatement in recent decades. Consequently, species discovery coupled with more quantitative assessments of distinct populations for reproductive isolation may help determine whether undescribed cryptic species might contribute some degree of overestimation to times required for the completion of speciation.

Our estimate of species diversification is largely consistent with a constant birth rate model with a net diversification rate of 0.0191 – 0.0248 per lineage per 10 million generations. Nonetheless, inspection of the lineage-through-time plot suggests a transient elevation in net diversification starting approximately 1 billion generations ago (Figure 3, Supplementary Figure S5). The confidence intervals are wide for the date estimates of these deep internal nodes of the *Caenorhabditis* phylogeny, however, and also show substantial gene tree discordance with the species tree (Figure 1). It is therefore possible that the hump in the LTT curve starting at ∼1 billion generations ago reflects spurious clustering in time of several deep splits in the phylogeny that are associated with long branches to small clades with relatively sparse numbers of known species.

For an assumption of 20 generations per year, there is a tantalizing coincidence of *Caenorhabditis* emerging as a genus following the radiations of terrestrial life in the middle of the Cretaceous period ∼100 Mya, and of angiosperms and insect pollinators in particular [85–87]. Given the common microenvironments inhabited by *Caenorhabditis* of microbe-rich rotting fruit, flowers, and vegetation and their phoretic associations with insects and other invertebrates [88, 89], further analysis of such deep phylogenetic coevolution may be warranted. Discovery and inclusion of additional basally-branching species of *Caenorhabditis* will help to better establish the timing of these ancestral nodes in the phylogeny, and also motivate statistically robust formal tests of co-diversification with other groups of organisms and abiotic variables related to climate and tectonic plate geography.

### Challenges to date estimation in units of years for *Caenorhabditis*

The conversion of divergence time into units of years for *Caenorhabditis* requires strong assumption about the effective number of generations per year in this group. We propose annualized dates based on 20 generations per year (Xu Wei and Marie-Anne Felix, personal communication), though the corresponding calculated date of 74 Mya root of the genus would double or halve if one were to instead assume 10 or 40 generations per year (i.e., shifting a Late Cretaceous root to one in the Late Jurassic or Eocene). By point of comparison, recent analysis of the phylum Nematoda that incorporates fossil calibrations from other nematode groups suggests common ancestry of *C. elegans* and *C. remanei* at ∼59 Mya, of *Caenorhabditis* and sister genus *Diploscapter* in the vicinity of 140-190 Mya, and for *Caenorhabditis* and *Pristionchus* at ∼290 Mya [90, 91]. Fossil evidence of Rhabditid nematodes generally, as well as their Diplogastrid relatives that includes *Pristionchus* [92], dates at least to the Cretaceous >66 Mya [93]; direct fossil evidence of nematodes dates to approximately 400 Mya [94] with the split with Arthropoda estimated at approximately 600 Mya [71]. While we lack fossil evidence for *Caenorhabditis* itself, even our deepest estimated timeframe for the duration of the genus is plausible given fossil evidence from other nematodes, for example, as for what appear to be congeners of extant *Heleidomermis* nematodes that occur in Burmese fossil amber from ∼100 Mya [93].

Several other factors contribute uncertainty to our estimates of time to common ancestry. First, our conversion of time in units of generations to years also presumes equivalent generation times among species. The possibility of variation in generation time – due perhaps to faster generation turnover in species that inhabit warm geographic regions with abundant food resources relative to species inhabiting cooler or food-scarce habitats or with highly specialized life histories – might present as molecular evolutionary rate heterogeneity among branches in the phylogeny and contribute a source of uncertainty in the date estimates [95, 96]. While some branches of the phylogeny deviated from the average rate of molecular evolution (Figure 1, Supplementary Figure S4), the relative differences were modest. Although some *Caenorhabditis* species are capable of passaging through >150 generations in a year under ideal laboratory conditions, the realized generation turnover will be restricted in nature by suboptimal abiotic and biotic circumstances as well as the incidence and duration of time spent in the quiescent dauer stage [88, 89]. Dauer quiescence may persist for the equivalent duration of several dozen generations of direct development [97]. To the extent that dauer larvae contribute a “seed bank” to population dynamics [98], they would act to extend the effective generation time of the species even beyond just the simple delay in onset of reproduction (and also contribute to elevated genetic effective population size) [99–101]. Future studies that document *Caenorhabditis* generation times in nature will help to refine divergence time estimates in units of years.

Our dating estimates also depend on mutation rate estimates from *C. elegans* being applicable across the genus. Mutation rates for *C. elegans* and *C. briggsae* and *P. pacificus* do appear to be largely consistent [102, 103], but it remains possible that other species experience different rates of mutation input [104]. For example, an alternative point estimate for the base substitution rate in *C. elegans* is ∼20% lower than the value used in our calibrations [103], the use of which would correspondingly lead to ∼20% more ancient divergence time estimates. Moreover, the broad range of effective population size (*N*_e_) among species of *Caenorhabditis* also suggests the possibility that variation in the efficacy of selection (*N*_e_*s*) among species, i.e. variation among lineages in the proportion of substitutions that fix by selection versus genetic drift, also could contribute a source of substitution rate heterogeneity in amino acid sequences [96]. In addition, we did not adjust dates for ancestral polymorphism (θ), so dates may overestimate divergence times by θ/(2µ) generations [67]. Among gonochoristic *Caenorhabditis*, known silent-site neutral θ varies from 0.01 to 0.14 [68], implying that dates may be overestimated by 2 to 30 million generations depending on the species. Future work that quantifies *N*_e_, µ, and generation times across the phylogeny can help to improve ancestral date estimates as well as our understanding of divergence in fundamental evolutionary properties.

### A collection of “speciation clocks” for *Caenorhabditis*

Since the pioneering analysis of Coyne and Orr [1] that first documented “speciation clocks” in *Drosophila*, speciation biologists have aimed to quantify the accumulation of distinct components of reproductive isolation as a function of time in a range of vertebrate, insect, plant, and fungal taxa [8, 12]. We present with *Caenorhabditis* the first set of reproductive isolation clocks for the nematode phylum. Because test-crosses form part of the best practices in species diagnosis for this group [36], we collated the abundant qualitative information from existing literature about the potential for interspecies mating, zygote formation, and production of hybrid offspring. We observed that post-zygotic isolation generally accrues prior to pre-zygotic isolation in *Caenorhabditis*, consistent with speciation in this group being driven by intrinsic barriers [11]. A caveat to this conclusion, however, is that *Caenorhabditis* niche space differences and the potential for extrinsic ecology-dependent reproductive isolation remain largely uncharacterized despite the accelerating ecological research on these organisms [88, 89, 105]. Nonetheless, because prezygotic reproductive isolation associates with ecological divergence in both animals and plants [106], a testable prediction for future research in *Caenorhabditis* is that species with stronger barriers to mating (e.g., *C. inopinata*, *C. portoensis*) may also occupy more distinctive ecological niches.

One caution in “speciation clock” inference is to guard against the presumption that reproductive isolation must inevitably increase through time, as speciation reversal is a possible future fate of species that are incompletely isolated [107–109]. Nonetheless, time-dependent reproductive isolation pervades taxa [1, 2, 9, 12, 19] and “speciation clocks” sometimes are argued to be consistent across disparate groups [27, 110]. In *Caenorhabditis*, reproductive isolation is complete in nearly all species pairs documented to date, making speciation reversal unlikely to influence the trends that we have demonstrated.

Broad taxonomic surveys have shown that diverse taxa show consistent molecular evidence of speciation when synonymous site divergence (*K*_S_) exceeds ∼5.5% [73], or approximately 2 My of divergence [27]. There exists a wide range of variability, however, with partial compatibility often extending up to 10 My or more in many organisms [2, 6]. It is possible that the high karyotype stability of holocentric chromosomes within *Caenorhabditis*, albeit with substantial gene rearrangement within chromosomes [111, 112], might facilitate incomplete isolation at even longer timescales as we have summarized here. Moreover, variability among species pairs within reproductive isolation clocks will partly reflect stochasticity in the contributions of lineage-specific selection and of large-effect incompatibilities relative to small-effect incompatibilities [2, 9, 113].

A limitation of our ability to time the accumulation of reproductive isolation in *Caenorhabditis* is that our analysis includes few cases of intermediate post-zygotic reproductive isolation. With the rapid pace of species discovery such that known species in culture has doubled in each of the last two decades, additional examples of incomplete isolation between distinct biological species are sure to present themselves [61]. For example, there may be a complex of cryptic species related to *C. remanei* in addition to the “previously hidden” species *C. latens*, given observed patterns of cross-compatibility [51]; the short genetic distances separating species related to *C*. *agridulce* remain to be fully documented for their magnitude of reproductive isolation [62]; intra-species phylogeographic splits and incompatibilities might warrant consideration of incipient speciation, even in the absence of splitting of formal species designations [114–116]; and newly-discovered species in the Sinica subclade show incomplete reproductive isolation [61].

More comprehensive future quantification of pre-mating isolation among *Caenorhabditis*, as well as distinguishing pre-mating from post-mating pre-zygotic isolation, also will help evaluate how robust is the macroevolutionary pattern of faster evolution of post-zygotic relative to pre-zygotic isolating barriers. Likewise, deeper investigation of the developmental timing at which reproductive isolation manifests can provide insight into possible developmental genetic rules in the evolution of post-zygotic barriers [22, 23]. Our observation that hybrid sterility evolves sooner than hybrid inviability, as documented in a variety of other taxa [18, 117], is consistent with the disproportionate fragility of developmental programs governing gonad development and similar to some logic invoked for the “faster male” and “fragile male” hypotheses to explain Haldane’s rule [118–120]. Incorporation of additional closely-related *Caenorhabditis* species pairs with less-extreme positions on the speciation continuum, and more refined quantitative metrics, will provide finer resolution and tests of how distinct isolating barriers accumulate in species formation and maintenance.

## Methods

### Species tree topology and orthogroup inference

We retrieved reference genome and transcriptome assemblies for 51 *Caenorhabditis* species (Supplementary Table S1) from a combination of WormBase ParaSite [121, 122], NCBI GenBank [123], and the Caenorhabditis Genomes Project [124], as described in [125]. For all 51 species, we used BUSCO v5.2.2 [126] on the set of annotated protein sequences (using the longest isoform for each gene) to detect the single-copy genes in the nematoda_odb10 gene set, then aligned them using PRANK v.170427 [127] to yield a total of 3130 alignments. We then constructed 3130 gene trees with IQ-TREE v2.3.6 [128], selecting the best model using ModelFinder [129], which were input into ASTRAL-Pro3 v1.19.3.6 [130] to infer a single species tree for which site concordance factors were calculated using IQ-TREE v2.3.6 [128]. We considered several possible alternate topologies for certain analyses below, but treated this ASTRAL species tree as the primary phylogenetic hypothesis for most subsequent analysis. Using the ASTRAL species tree topology, we then defined groups of orthologous genes (orthogroups) between all 51 *Caenorhabditis* species with OrthoFinder v2.5.2 [131], as described in [125]. For each single-copy orthogroup (all 51 species having exactly one ortholog present), we aligned the DNA coding sequences with PRANK v.170427 [127] in codon-alignment mode and using GUIDANCE v2.0.2 [132] with 50 bootstrap runs and a cutoff score of 0.93 to remove codons that could not be reliably aligned. This resulted in a set of 205 aligned single-copy orthogroups each with representation from all 51 species. Supplementary data files and analysis scripts available at https://github.com/Cutterlab/Caenorhabditis_Divergence_Times_Speciation_Clocks.

### Calibration node selection and time calculation

Because *Caenorhabditis* lack fossil evidence for timetree calibration, we exploited experimental determinations of neutral mutational input to calibrate molecular divergence of the phylogeny as a whole [63, 66], harnessing the most recently-diverged species pairs (10 distinct speciation events). We identified the 11 most recently-diverged species pairs as those with the lowest *K*_S_ values (i.e. unadjusted *K*_S_ < 0.4) from calculations with MEGA v11.0.10 of Jukes-Cantor corrected synonymous substitution rate (*K*_S_) between every pair of species based on the concatenation of the 205 single-copy orthogroup coding sequence alignments [133] (Supplementary Figure S1A).

Under the molecular clock hypothesis [134], neutral divergence accumulates linearly with time (*T*) and synonymous-site change typically best reflects this evolutionary behavior (i.e., *K*_S_ = 2µ*T* for pairwise neutral divergence *K*_S_ and neutral mutation rate µ). However, high divergence leading to saturation of synonymous-site substitutions, as well as selection for biased codon usage among highly-expressed genes in *Caenorhabditis*, compromises this assumption without appropriate accounting [135, 136]. We individually aligned the coding sequences of all 1:1 orthologs between each of the 11 species pairs with PRANK v.170427 [127] in codon-alignment mode. We then estimated *K*_S_ for each 1:1 ortholog alignment using the FitMG94 script from HyPhy v2.5.52 [137] after replacing all stop codons with gaps, then excluded those with *K*_S_ > 1 to yield from 8774 to 15,204 values per species pair. Additionally, the Effective Number of Codons (ENC) was calculated for each 1:1 ortholog alignment using coRdon v1.14.0 (https://github.com/BioinfoHR/coRdon), based on the first sequence in each alignment. After removing alignments with *K*_S_ > 1 (representing likely errors in sequence alignment or ortholog inference), we performed a linear correction for the effect of ENC on *K*_S_ for each species pair, as in [63]. Specifically, we calculated a corrected *K*_S_ value (*K*_S_′) for each 1:1 ortholog pair as *K*_S_′ = *K*_S_ + *a* * (61 – ENC), where *a* is the slope of the *K*_S_–ENC regression line for that species pair, and 61 is the value of ENC representing no codon usage bias.

The median *K*_S_′ values for these 11 species pairs based on >8,000 genes per species pair thus represent our best estimates of neutral divergence between *Caenorhabditis* species that do not experience saturation of substitutions, and account for multiple mutational hits and the effects of selection on synonymous sites. The *K*_S_′ values were then converted to divergence times using the molecular clock formula *K*_S_′ = 2µ*T*, where *T* is the divergence time (in generations) and µ is the mutation rate. We applied the neutral mutation rate of µ = 2.33 x 10^-9^ mutations per site per generation from experimental estimates of the base-substitution mutation rate from genome sequencing of long-term mutation accumulation studies in *C. elegans* [66]. We chose to calibrate the *C.* sp. 8 – *C.* sp. 56 – *C. agridulce* ancestral node of the phylogeny using just the parameters for the *C.* sp. 8 – *C.* sp. 56 pair. Supplementary data files and analysis scripts available at https://github.com/Cutterlab/Caenorhabditis_Divergence_Times_Speciation_Clocks.

### Date estimation for the *Caenorhabditis* phylogeny

The extensive divergence due to deep ancestry of many *Caenorhabditis* species renders synonymous-site divergence saturated with substitutions (i.e. *K*_S_ > 1), which erases information content about linear neutral mutation accumulation over time for such sites (Supplementary Figure S1B). While corrections like we made in calculating *K*_S_′ attempts to counter this effect, sufficiently large divergence nonetheless is expected to lead to the compression of deep branch length estimation and a corresponding violation of the presumption of rate constancy [138]. We therefore incorporated amino acid sequence divergence to date the *Caenorhabditis* phylogeny for all 51 species, expanding on the 10 *K*_S_′-derived calibration nodes described above, using the StarBeast3 v1.1.8 package of BEAST 2 v2.7.5 [139, 140].

A single run of BEAST included all 205 single-copy ortholog alignments as separate partitions, with each alignment using its best-fit sequence evolution model as determined by ModelFinder in IQ-TREE v2.1.2 [128, 129]. We used the ASTRAL tree topology with a relaxed clock model, specifying clock rate priors of 1/X for each alignment, and providing our inferred distributions of calibration times as priors for the 10 calibration nodes. Divergence times for the calibration node priors provided to BEAST were converted to units of millions of years assuming 10 generations per year, defining priors based on gamma distribution parameter fits to the distributions of calculated divergence times for each species pair using the fitdistr() function in R v4.2.1 [141] (except the *C. drosophilae* – *C.* sp. 2 pair, for which a log-normal distribution better fit the data based on log-likelihood) (Supplementary Figure S2). Note that posterior date estimates output from BEAST can then be linearly scaled to alternative assumptions about generation times per year. The chain was run for 200,000,000 steps (90,000,000 step burn-in), sampled every 5000 steps, and convergence was assessed using Tracer v1.7.2 [142]. Divergence times were taken to be the mean of the associated posterior distribution after burn-in. These improved approaches to handling mutational saturation between orthologs, as well as updated estimates of µ and more extensive taxon sampling, influence the date estimates relative to those reported previously [63]. To estimate divergence time in years, we presume all *Caenorhabditis* species to experience identical effective generation times [143] from a range of 10 to 40 generations per year [63](Xu Wei and Marie-Anne Felix, personal communication).

We then created a lineage-through-time plot based on this dated phylogeny using the R package paleotree v3.4.7 [144]. To compare this plot to the expectation under a pure-birth process, we used phytools v2.4.4 [145] to calculate Pybus & Harvey’s γ statistic [84], and estimated the diversification rate for our phylogeny using the fit.bd() function assuming species sampling fractions of either 100% or 59% (i.e. 51/86, based on the 86 total *Caenorhabditis* species known to date [61]). For each of these estimated rates, we determined the average lineage-through-time plot from 1000 phylogenies simulated with that speciation rate (and a death rate of 0) and the same total age as our primary phylogeny, either allowing the simulated trees to have any final number of tips (for the sampling fraction of 59%) or requiring all simulated trees to have 51 tips (i.e sampling fraction of 100%).

We assessed the robustness of time estimates in three ways. First, to assess sensitivity of divergence time estimates to gene composition, we split the set of 205 single-copy ortholog alignments into 4 arbitrary subsets of 51 or 52 alignments each, and reran BEAST on each alignment subset as described above. BEAST chains for these subsets were run for 25,000,000 steps each, sampling every 2500 steps and using a burn-in of either 7,500,000 steps (for subsets 1 and 4) or 9,000,000 steps (for subsets 2 and 3). Second, to assess sensitivity of divergence time estimates to species tree topology, we inferred four alternate species tree topologies from modifications to our tree inference pipeline. These alternate trees that exhibited minor differences in topology were obtained by: (1) filtering our BUSCO protein sequence alignments with GUIDANCE v2.0.2 [132] using 50 bootstrap runs and a cutoff score of 0.93, followed by inferring gene trees with IQ-TREE (for a total of 1768 gene trees) and the species tree with ASTRAL-Pro3; (2) inferring the species tree with IQ-TREE after concatenating all 3130 BUSCO protein sequence alignments into a single alignment; (3) using PRANK to align the protein sequences of any orthogroups with at least 40 species represented (allowing for multicopy genes rather than only single-copy BUSCOs), followed by inferring gene trees with IQ-TREE (for a total of 9926 gene trees) and the species tree with ASTRAL-Pro3 [146–148]; and (4) allowing BEAST to infer its own species tree topology simultaneously with the divergence times, instead of providing it a fixed tree topology (aside from the 10 calibration points, which remained fixed). Each of these alternate tree topologies (indicated in Figure 1) was used to re-estimate divergence times with BEAST, as above, using the 51 alignments in gene subset 3. Chains for these four runs of BEAST were run for 25,000,000 steps each, sampling every 2500 steps and using a burn-in of either 12,000,000 steps (for the BEAST-inferred tree topology) or 9,000,000 steps (for the other topologies).

Finally, to assess the sensitivity of divergence time estimates to inclusion of basally-branching taxa, many of which have long terminal branches, we carried out our dating procedure restricted to the 31 species of the Elegans supergroup. We repeated our pipeline of aligning DNA coding sequences of single-copy orthologs with PRANK and GUIDANCE followed by divergence time estimation with BEAST, using a larger set of 692 orthogroups that were single-copy in the 31 species belonging to the Elegans supergroup (orthogroup sequences from the remaining 20 species were excluded). These 692 orthogroups were divided into four subsets: (1) the 205 orthogroups included in our original 51-species run of BEAST, and (2-4) three subsets of 162 or 163 arbitrarily-selected orthogroups from the remaining 487 orthogroups. We ran BEAST on these four subsets as above, specifying the topology of the Elegans supergroup based on the ASTRAL species tree (thus only including calibration time priors for the 5 calibration species pairs within the Elegans supergroup). For the 205-orthogroup run, we used a chain length of 150,000,000 (80,000,000 step burn-in), sampling every 5000 steps. For the remaining three runs, we used a chain length of 100,000,000 (60,000,000 step burn-in), sampling every 5000 steps. Supplementary data files and analysis scripts available at https://github.com/Cutterlab/Caenorhabditis_Divergence_Times_Speciation_Clocks.

### Extending the *Caenorhabditis* timetree to inclusion of new species

To link the BEAST-derived estimates of divergence time (*T*) with protein divergence more generally, we established the relation between non-synonymous site substitution rates (*K*_A_) for all 1:1 orthologs between a pair of species and *T*. This approach allows an approximated inference of *T* for any new nodes in the tree that arise from consideration of additional species, from *K*_A_ alone, without the time-intensive need to rerun BEAST on all species. For every unique pairwise combination of our 51 species, we aligned the DNA coding sequences for all 1:1 orthologs between these species using PRANK v.170427 [127] in codon-alignment mode and calculated the non-synonymous substitution rate per site (*K*_A_) as well as *K*_S_ for each resulting alignment using the FitMG94 script from HyPhy v2.5.52 [137] after replacing all stop codons with gaps. We then computed the median *K*_A_ value and median *K*_S_ value across all aligned 1:1 orthologs for each species pair, averaging across 4390 to 15,341 values per species pair. For each node on the species tree, we then calculated the median across corresponding species-pairs of their ortholog median *K*_A_ values and regressed them linearly against BEAST estimates of divergence time to obtain a regression equation relating *K*_A_ and *T*.

As test cases of this approach, we obtained genome assemblies for *C. auriculariae* and *C. niphades* from the Caenorhabditis Genomes Project [124] and NCBI GenBank accession GCA_946814055.1 [65, 123], respectively. To run OrthoFinder, we determined the phylogenetic placement of *C. niphades* by repeating our single-copy BUSCO-based ASTRAL species tree inference pipeline, but only using *C. niphades*, the 12 species in the Japonica group, and *C. elegans* and *C. briggsae* as outgroups (Supplementary Figure S3). *Caenorhabditis auriculariae* was taken to be sister to *C. monodelphis* [64, 65]. We then reran OrthoFinder v2.5.2 [131] to also include *C. auriculariae* and *C. niphades*, aligned 1:1 orthologs, and calculated median *K*_A_ values between *C. auriculariae*, *C. niphades* and the other 51 species, as described above. Estimates of divergence times for *C. auriculariae* and *C. niphades* based on *K*_A_ with other species were then computed from the regression equation for *K*_A_ and *T*. Supplementary data files and analysis scripts available at https://github.com/Cutterlab/Caenorhabditis_Divergence_Times_Speciation_Clocks.

### Reproductive isolation inference

We compiled published accounts of cross compatibility for 258 *Caenorhabditis* inter-species pairings and 5 species with pairings of distinct populations within a species (https://github.com/Cutterlab/Caenorhabditis_Divergence_Times_Speciation_Clocks). We defined overall F1 reproductive isolation (*R_F1_*) from 0 (production of fully viable and fertile adults) to 1 (no fertile adults produced), with intermediate values when possible calculated as 1-(intercross fitness)/(intracross fitness) as for outbreeding depression calculations in [149] and the *RI*_1_ metric of [150]. We defined a similar metric for F2 reproductive isolation (*R_F2_*), setting values to 1 if the F1 value was 1. A pre-mating isolation metric (*R_PreM_*) ranged from 0 (copulation occurs) to 1 (no copulation), with values of 0 also inferred indirectly from the production of hybrid embryos or later stages; values of 0.5 were assigned in cases where copulation was reported with no sperm transfer [70], although such instances likely are under-reported. Pre-zygotic isolation (*R_PreZ_*) included cases with evidence of either pre-mating isolation or post-mating pre-zygotic barriers (e.g., production of unfertilized oocytes but no embryos), with the metric also ranging from 0 to 1; values of 0.5 correspond to cases of ectopic sperm invasion as a form of gametic isolation [57]. Data relevant to the *R_PreZ_* metric also is subject to under-reporting in the existing literature on *Caenorhabditis*. We also defined a metric for post-zygotic isolation (*R_PostZ_*) ranging from 0 to 1 for the subset of cases where fertilized embryos or later stages of hybrid development were reported (0=fully fertile adults, 1=all sterile or inviable F1, intermediate values according to the above outbreeding depression equation when quantitative reports allowed). Intermediate values for these metrics were quantified in only a minority of cases in the literature, and so most cross compatibility scores of 0 or 1 were allocated based on the presence of any hybrids fitting the metric criteria. Multiple reported values for conspecifics were averaged in analysis. Finally, we categorized the latest stage of F1 development achieved in a hybrid cross (fertile adult, sterile adult, larvae, embryogenesis, ambiguous reporting as to whether the cross yielded fertilized embryos or reported unfertilized oocytes/failure to copulate). We then tested for an association of these ordinal categories with divergence time using Spearman rank correlation with cor.test() in R.

For each metric of reproductive isolation (*R_i_*), we fit a two-parameter logistic function using the nls() function in R with respect to divergence time in generations (*T*) to estimate the model coefficients *a* and *b* according to *R_i_* = 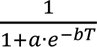 (where *b* represents the rate of accumulation of reproductive isolation barriers and *a* is proportional to the difference in fitness within versus between species) [18, 151]. The expected time to 50% reproductive isolation is thus *T*_*R_i_*=0.5_ = 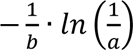) and the expected time to 95% reproductive isolation is *T*_*R_i_*=0.95_ = 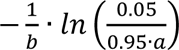. Note that shared phylogenetic history makes points not fully independent in these plots based on all species pairs. To account for phylogeny, we used a greedy algorithm to select the maximum number of phylogenetically independent species pairs with measurements of each type of reproductive isolation. This procedure yielded 12 to 16 species pairs per metric, for which we then averaged values from reciprocal cross directions for a given species pair and repeated the logistic function fits. Supplementary data files and analysis scripts available at https://github.com/Cutterlab/Caenorhabditis_Divergence_Times_Speciation_Clocks.

## Data availability

Reference genome and transcriptome assemblies for 51 *Caenorhabditis* species were retrieved from a combination of WormBase ParaSite [121, 122], NCBI GenBank [123], and the Caenorhabditis Genomes Project [124] and listed in Supplementary Table S1. Data files and analysis scripts available on GitHub at https://github.com/Cutterlab/Caenorhabditis_Divergence_Times_Speciation_Clocks.

## Acknowledgements

We are grateful to Lewis Stevens and Mark Blaxter of the Wellcome-Sanger Institute for sharing *Caenorhabditis* genome data, to Xu Wei and Marie-Anne Felix for sharing estimates of generation time, Joshua Dall’Acqua for assistance in constructing an algorithm for selecting the largest number of phylogenetic contrasts, and to the *Caenorhabditis* evolution research community for generating data on reproductive isolation in the course of diagnosing new species. A.D.C. is supported by Discovery Grant funds from the Natural Sciences and Engineering Research Council (NSERC) of Canada (RGPIN-2018-05098); D.D.F. is supported by an NSERC Postgraduate Doctoral scholarship.

## Supplementary Tables

**Supplementary Table S1:** Sources of the genome/transcriptome assemblies and gene annotation files for the 51 *Caenorhabditis* species used in our primary divergence time estimates, as well as *C. auriculariae* and *C. niphades*. All data from the Caenorhabditis Genomes Project are also available at https://doi.org/10.5281/zenodo.12633738.

**Supplementary Table S2**: Phylogenetically-independent species pairs used to generated logistic fits for pre-zygotic, post-zygotic and F1 reproductive isolation and for regression analysis on latest viable stage of F1 hybrids.

**Supplementary Table S3:** For every pair of the 51 *Caenorhabditis* species used in our primary divergence time estimates, median K_A_ and median K_S_ across 1:1 orthologs, the number of 1:1 ortholog pairs used to calculate these medians, and our primary divergence time estimates based on 205 single-copy orthologs (i.e. times shown in Fig. 1, in units of millions of generations).

**Supplementary Table S4:** The values of K_S_ (both uncorrected and corrected, i.e. *K*_S_′) and ENC for every pair of 1:1 orthologs used to estimate calibration time priors for our 11 calibration species pairs (i.e. 1:1 ortholog pairs with uncorrected K_S_ ≤ 1).

**Supplementary Table S5:** For every species pair involving either *C. auriculariae* or *C. niphades* in comparison to one of our 51 primary species (or to each other), median K_A_ and median K_S_ across 1:1 orthologs, the number of 1:1 ortholog pairs used to calculate these medians, and the estimate of divergence time based on the median K_A_ value and our *K*_A_– divergence time regression.

**Supplementary Table S6:** Logistic function fits to components of reproductive isolation (RI) when conspecific divergence times are set to 0 or excluded.

## Supplementary Figures

**Figure S1.**
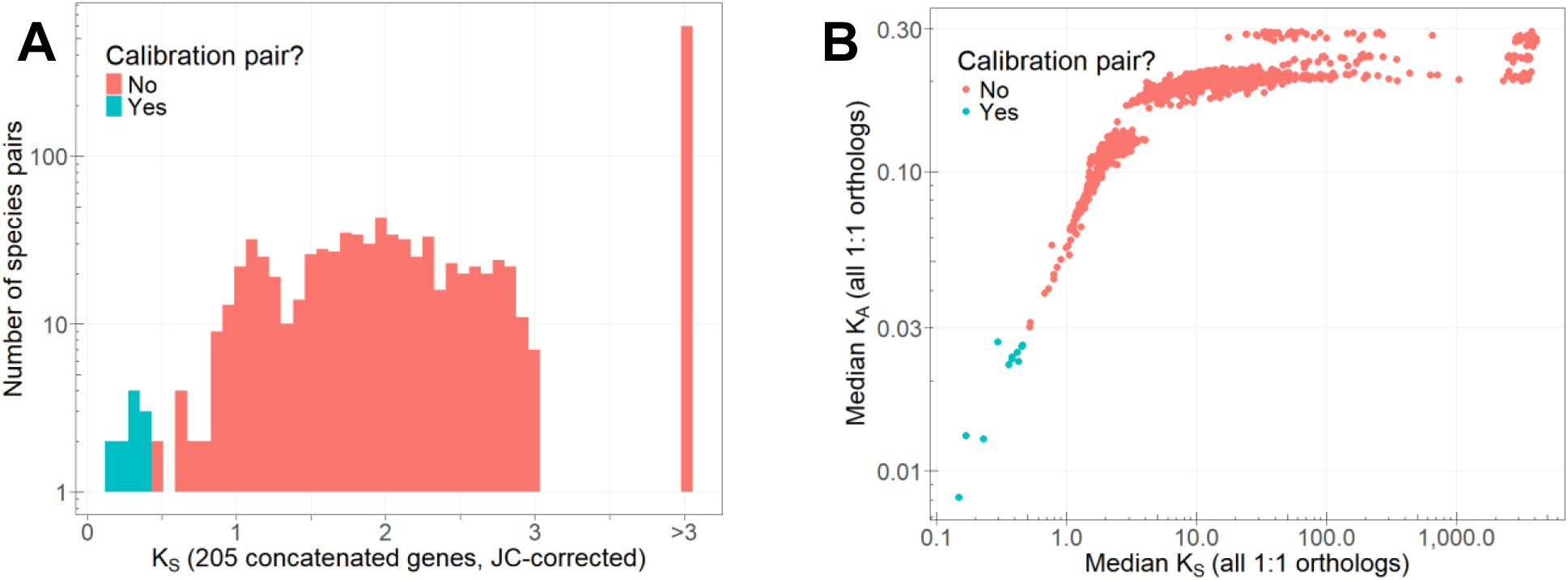
**A.** Histogram of Jukes-Cantor corrected values of synonymous-site substitution rate (*K*_S_) between all 1275 pairs of the 51 species used for divergence time estimation, calculated in MEGA based on the concatenation of the 205 single-copy orthologs. The 11 species pairs chosen for use in calibration include: *C.* sp. 56 – *C. agridulce*, *C. remanei* – *C. latens*, *C. nigoni* – *C. briggsae*, *C. drosophilae* – *C.* sp. 2, *C. brenneri* – *C.* sp. 48, *C. angaria* – *C. castelli*, *C. oiwi* – *C. kamaaina*, *C. dolens* – *C. quiockensis*, *C. becei* – *C. nouraguensis*, *C.* sp. 56 – *C.* sp. 8, and *C. agridulce* – *C.* sp. 8. **B.** Comparison of median *K*_S_ and median *K*_A_ between all 1275 pairs of the 51 species used for divergence time estimation, calculated with FitMG94 based on all 1:1 orthologs between each species pair.

**Figure S2.**
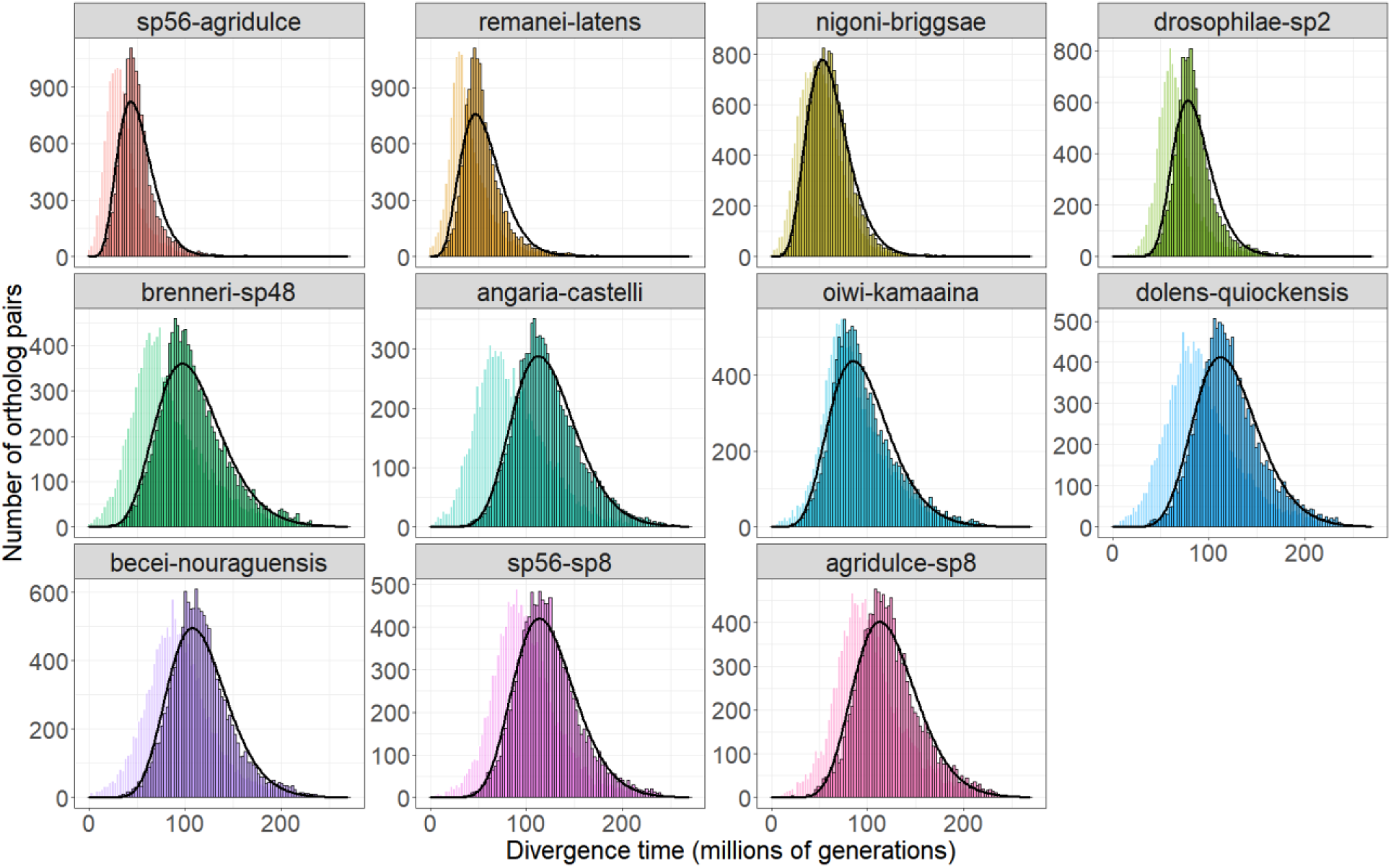
Distributions of divergence times for each calibration species pair, calculated by converting ENC-corrected values of *K*_S_′ (foreground, black-outlined bars) or uncorrected values of *K*_S_ (background, lighter bars) for 1:1 orthologs to divergence times using the strict molecular clock with an experimentally-derived estimate of mutation rate. Black curves on each panel show the density of the prior distribution that was given to BEAST for each species pair (for the common ancestor of *C.* sp. 8, *C.* sp. 56, and *C. agridulce*, only the *C.* sp. 8 – *C.* sp. 56 pair was used), based on fits to a gamma distribution (except the *C. drosophilae* – *C.* sp. 2 pair, for which a log-normal distribution better fit the data based on log-likelihood).

**Figure S3.**
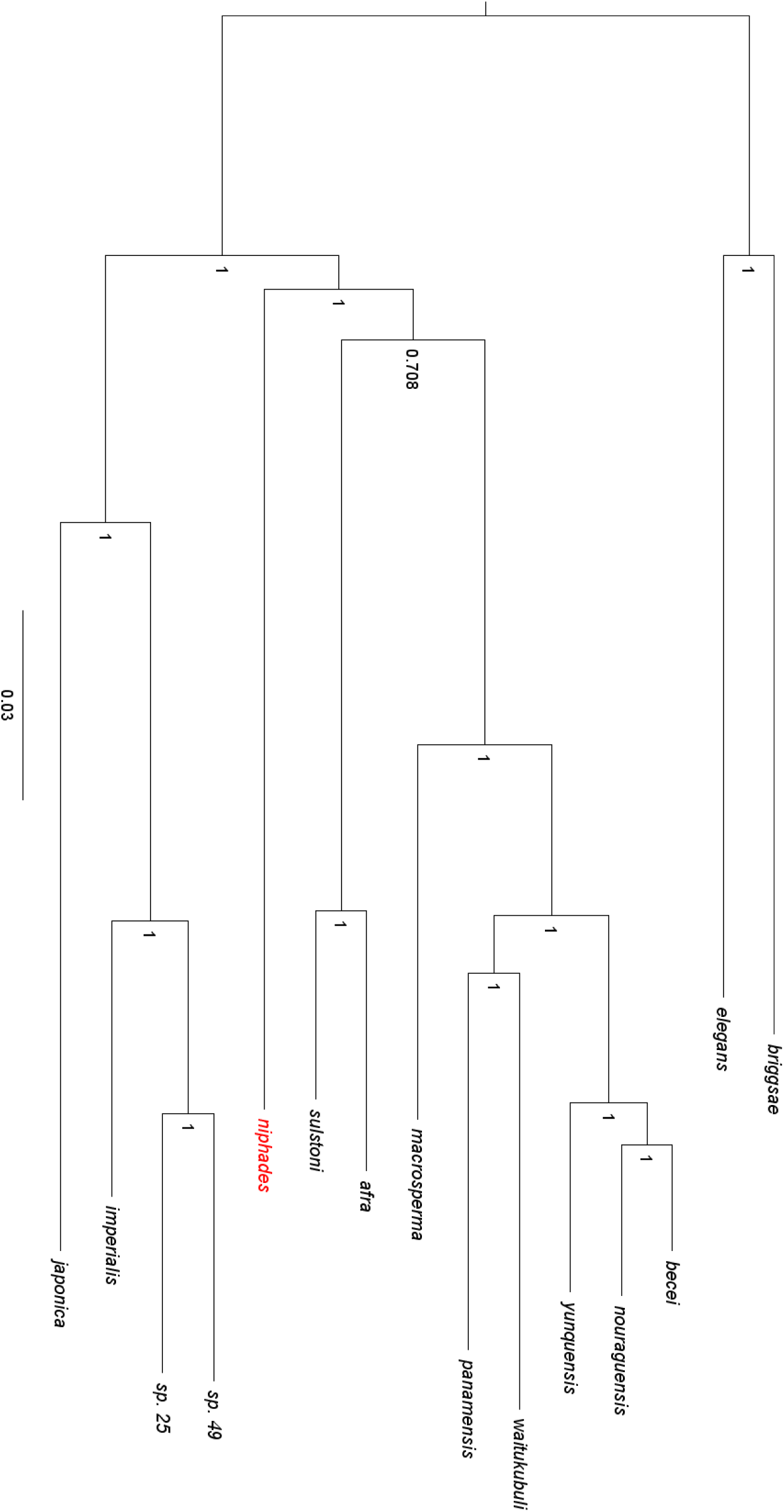
ASTRAL species tree of the Japonica group of species (including *C. niphades*, in red), as well as the outgroups *C. elegans* and *C. briggsae*. Node labels indicate posterior probabilities. Branch lengths are in units of amino acid substitutions per site.

**Figure S4.**
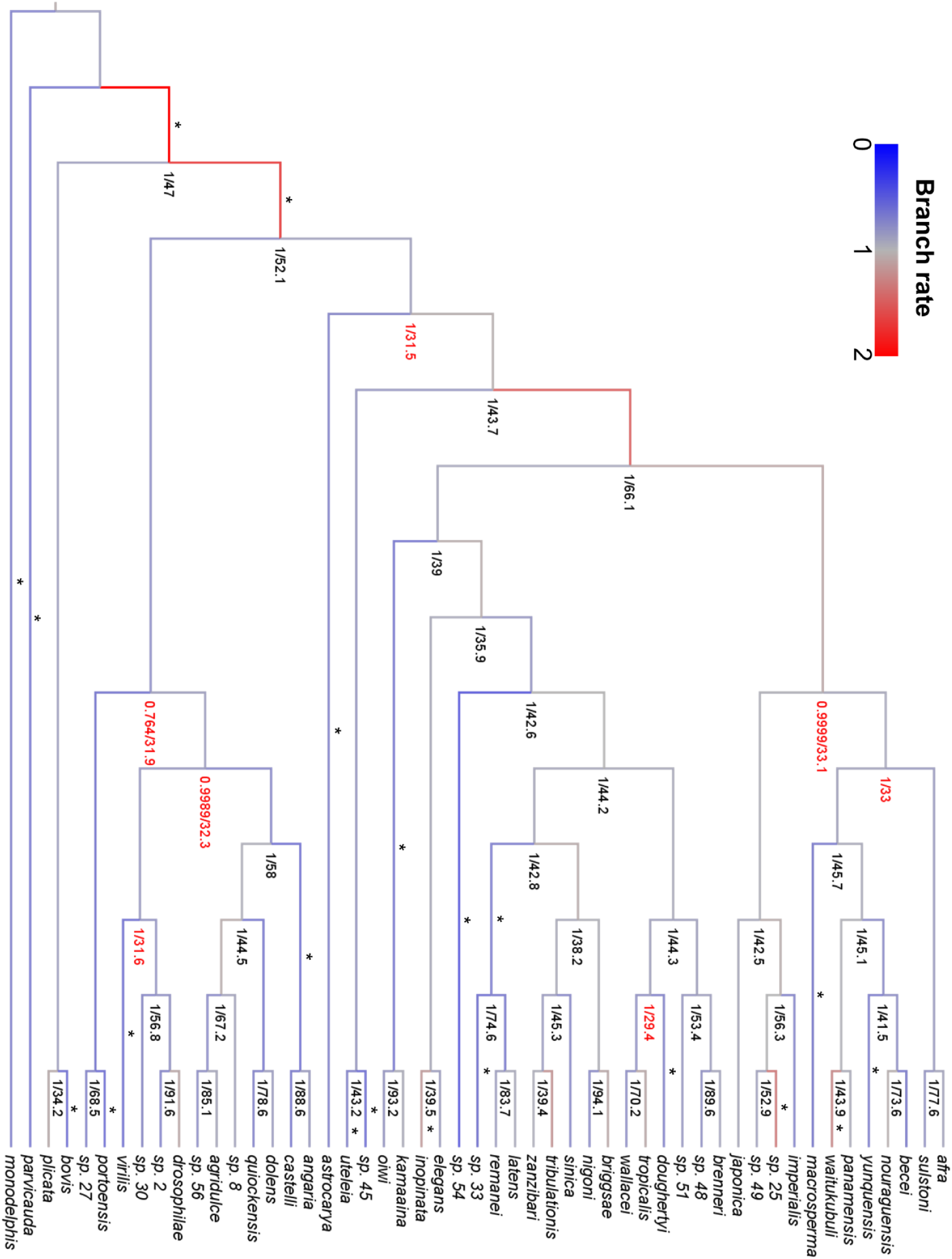
Topology of the ASTRAL species tree used for our primary divergence time estimates (Figure 1). Numerical labels on each node give the posterior probability (out of 1) followed by the site concordance factor (out of 100). Nodes with low statistical support (posterior probability < 1) or high discordance (site concordance factor < 33.3%) are labeled in red. The color of each branch gives its relative substitution rate under the relaxed clock model, with higher rates indicated by shades of red and slower rates indicated by shades of blue. Branches with a “*” have a 95% Highest Posterior Density interval for the estimated branch rate that does not overlap with 1.

**Figure S5.**
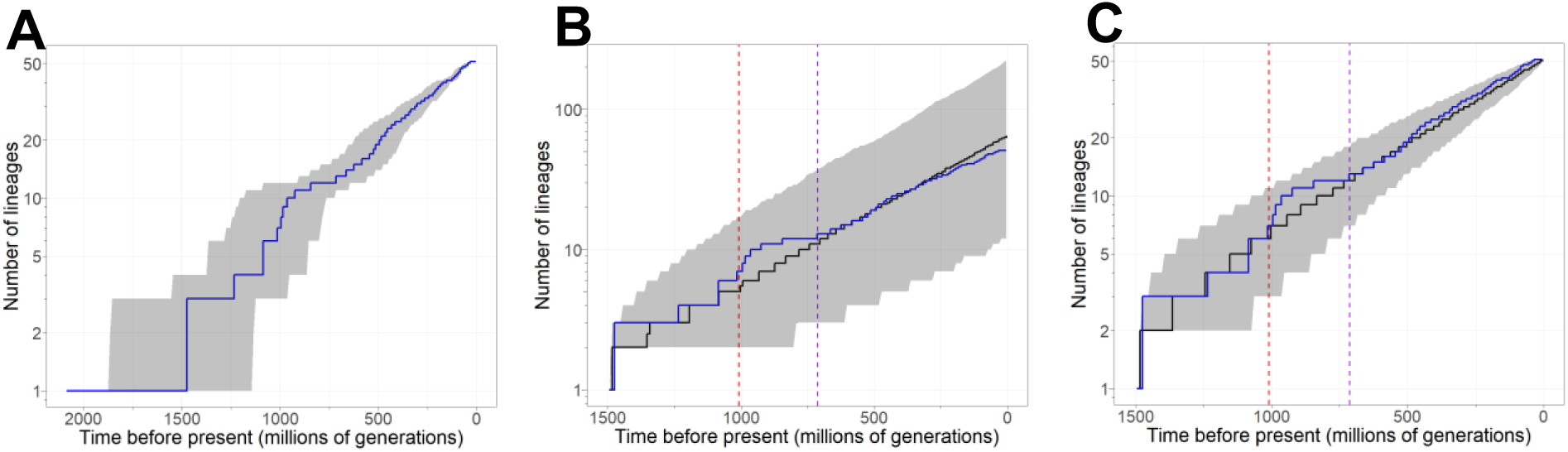
**A.** Lineage-through-time plot for the *Caenorhabditis* genus. Shaded intervals give the 95% upper and lower quantiles around the median, based on the posterior distributions of divergence times calculated by BEAST. Blue lines in all panels indicate the cumulative observed number of lineages through time for the 51-species phylogeny. **B.** Lineage-through-time plot for 1000 random phylogenies with the same total age as our primary phylogeny, but allowing any final number of tips given the estimated birth rate. Phylogenies were generated under the birth rate (λ=0.0248 per lineage per 10 million generations) estimated from our primary phylogeny when assuming a species sampling fraction of 59% (51/86). Shaded intervals give the 95% upper and lower quantiles around the median, based on the 1000 random trees. **C.** Lineage-through-time plot for 1000 random phylogenies, constrained to have the same total age and final number of tips (51) as our primary phylogeny. Phylogenies were generated under the birth rate (λ=0.0191) estimated from our primary phylogeny when assuming a species sampling fraction of 100%. Shaded intervals give the 95% upper and lower quantiles around the median, based on the 1000 random trees. Vertical dashed lines in B-C indicate the ages of the Elegans supergroup (purple) and the basal clade defined by the most recent common ancestor of *C. angaria* and *C. portoensis* (red).

**Figure S6.**
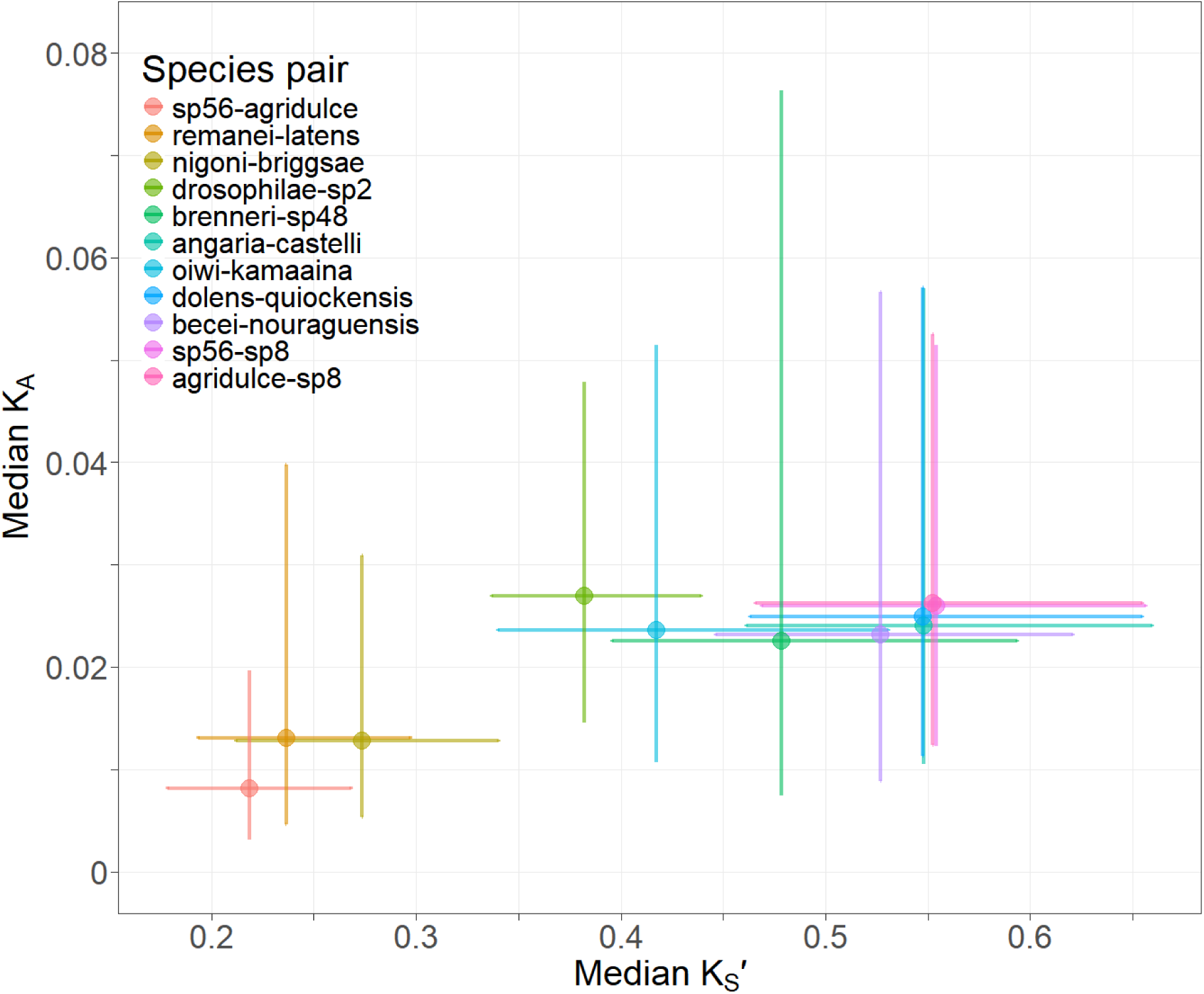
Median divergence across 1:1 orthologs for replacement site differences (*K*_A_) and synonymous site differences corrected for codon usage bias (*K*_S_′) for the 11 pairs of *Caenorhabditis* species used for divergence time calibration. Number of 1:1 ortholog pairs per species pair ranges from 9144 to 15,341 for *K*_A_ (median 13,020) and 8774 to 15,204 for *K*_S_′ (median 12,605). Error bars indicate interquartile ranges.

**Figure S7.**
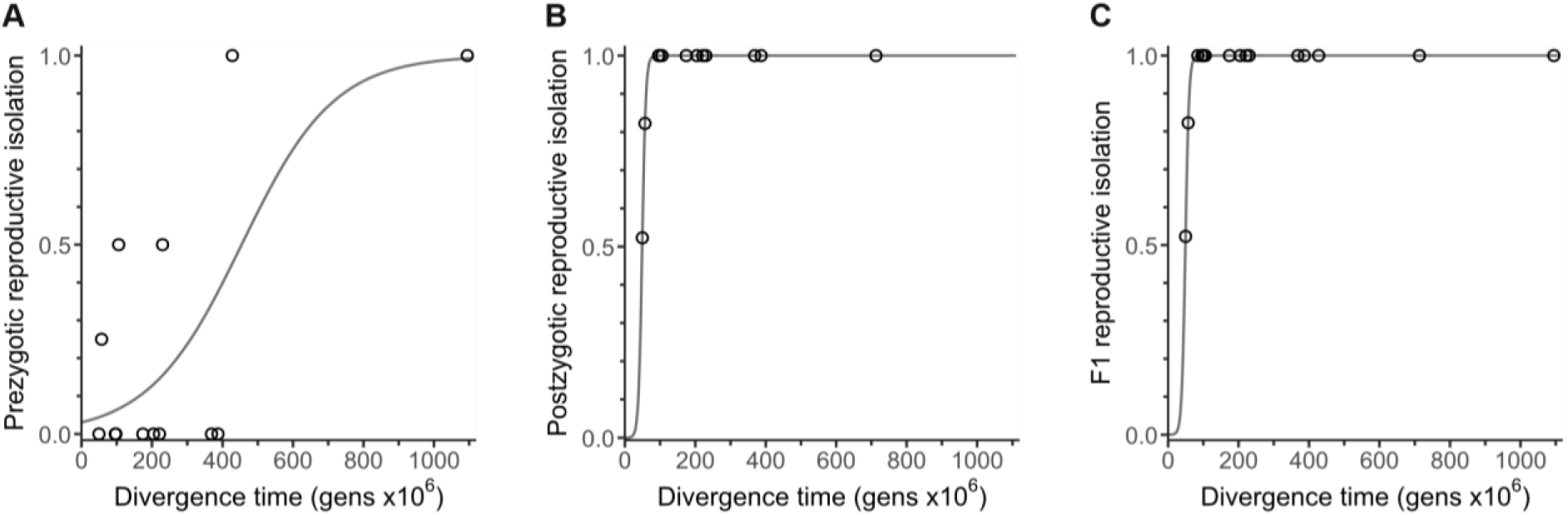
Reproductive isolation clocks generated using 16 phylogenetically independent species pairs (some RI metrics have fewer pairs), with reciprocal cross values averaged for a given pair.

**Figure S8.**
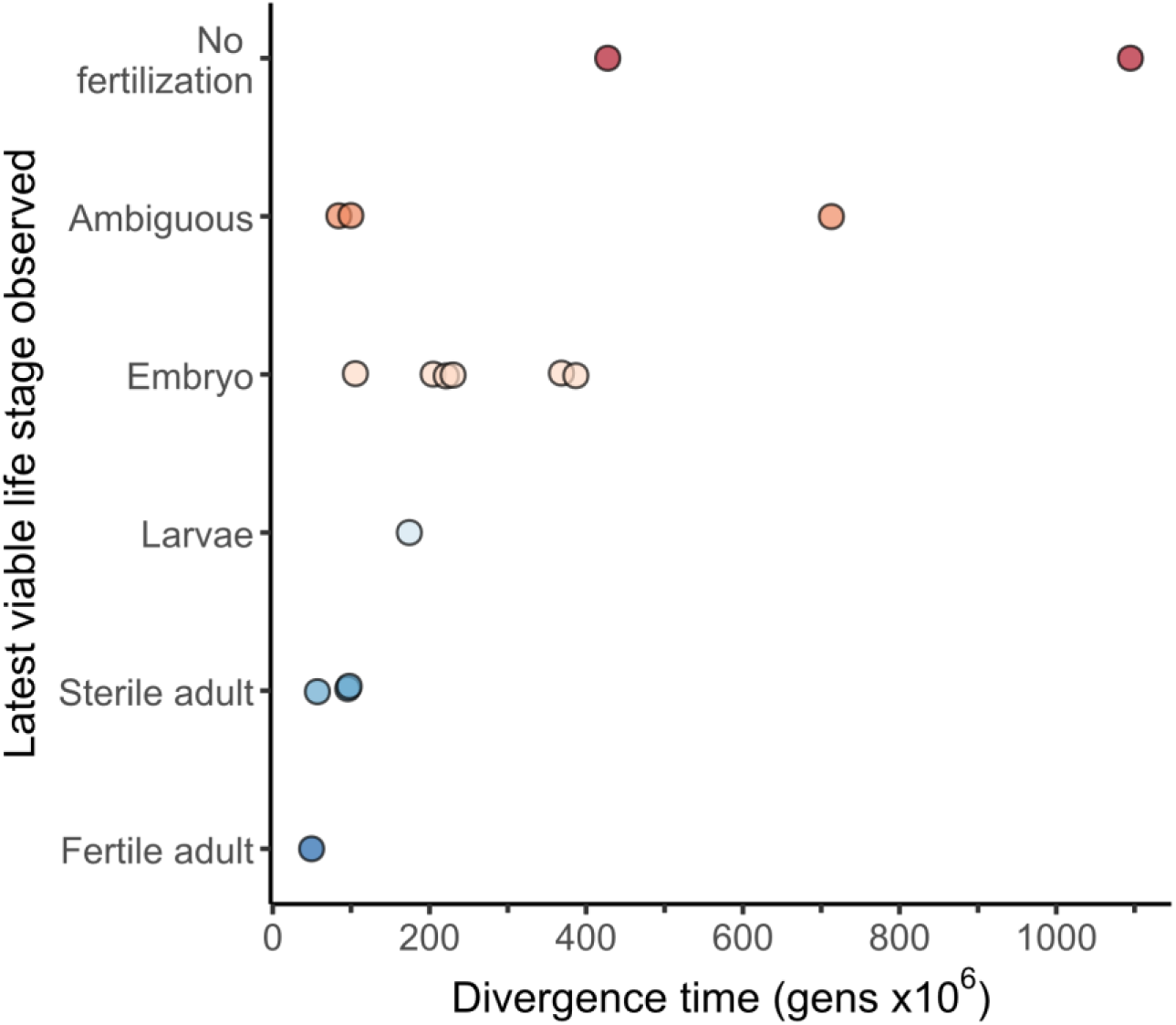
Latest stage reached by F1 offspring of 16 phylogenetically-independent species pairs. In cases where reciprocal cross directions produced F1 offspring that reached different terminal life stages, the latest stage is indicated.

## Literature cited

1. Coyne JA, Orr HA. Patterns of speciation in *Drosophila*. Evolution. 1989;43(2):362–81.

2. Edmands S. Does parental divergence predict reproductive compatibility? Trends Ecol Evol. 2002;17(11):520–7.

3. Coyne JA, Orr HA. Speciation. Sunderland, MA: Sinauer; 2004.

4. Rundle HD, Whitlock MC. A genetic inerpretation of ecologically dependent isolation. Evolution. 2001;55(1):198–201. doi: 10.1111/j.0014-3820.2001.tb01284.x.

5. Matute DR, Cooper BS. Comparative studies on speciation: 30 years since Coyne and Orr. Evolution. 2021;75(4):764–78. doi: 10.1111/evo.14181.

6. Bolnick DI, Near TJ. Tempo of hybrid inviability in centrarchid fishes (Teleostei: Centrarchidae). Evolution. 2005;59(8):1754–67. PubMed PMID: ISI:000231658900013.

7. Coyne JA, Orr HA. The evolutionary genetics of speciation. Philos Trans R Soc Lond B-Biol Sci. 1998;353(1366):287–305.

8. Giraud T, Gourbière S. The tempo and modes of evolution of reproductive isolation in fungi. Heredity. 2012;109(4):204–14. doi: 10.1038/hdy.2012.30.

9. Gourbiere S, Mallet J. Are species real? The shape of the species boundary with exponential failure, reinforcement, and the “missing snowball”. Evolution. 2010;64(1):1–24. doi: DOI 10.1111/j.1558-5646.2009.00844.x. PubMed PMID: ISI:000273455300001.

10. Mendelson TC, Inouye BD, Rausher MD. Quantifying patterns in the evolution of reproductive isolation. Evolution. 2004;58(7):1424–33. doi: 10.1111/j.0014-3820.2004.tb01724.x.

11. Seehausen O, Butlin RK, Keller I, Wagner CE, Boughman JW, Hohenlohe PA, et al. Genomics and the origin of species. Nat Rev Genet. 2014;15(3):176–92.

12. Coughlan JM, Matute DR. The importance of intrinsic postzygotic barriers throughout the speciation process. Philos Trans R Soc Lond B-Biol Sci. 2020;375(1806):20190533. doi: doi:10.1098/rstb.2019.0533.

13. Christianson SJ, Swallow JG, Wilkinson GS. Rapid evolution of postzygotic reproductive isolation in stalk-eyed flies. Evolution. 2005;59(4):849–57. doi: 10.1111/j.0014-3820.2005.tb01758.x.

14. Jewell C, Papineau AD, Freyre R, Moyle LC. Patterns of reproductive isolation in *Nolana* (Chilean bellflower). Evolution. 2012;66(8):2628–36. doi: 10.1111/j.1558-5646.2012.01607.x.

15. Palmer CA, Edmands S. Mate choice in the face of both inbreeding and outbreeding depression in the intertidal copepod Tigriopus californicus. Mar Biol. 2000;136(4):693–8. doi: 10.1007/s002270050729.

16. Kozak GM, Rudolph AB, Colon BL, Fuller RC. Postzygotic isolation evolves before prezygotic isolation between fresh and saltwater populations of the rainwater lillifish, *Lucania parva*. International Journal of Evolutionary Biology. 2012;2012:523967. doi: 10.1155/2012/523967.

17. Widmer A, Lexer C, Cozzolino S. Evolution of reproductive isolation in plants. Heredity. 2009;102(1):31–8. doi: 10.1038/hdy.2008.69.

18. Turissini DA, McGirr JA, Patel SS, David JR, Matute DR. The rate of evolution of postmating-prezygotic reproductive isolation in *Drosophila*. Mol Biol Evol. 2017;35(2):312–34. doi: 10.1093/molbev/msx271.

19. Coyne JA, Orr HA. “Patterns of speciation in *Drosophila*” revisited. Evolution. 1997;51(1):295–303.

20. Bundus JD, Alaei R, Cutter AD. Gametic selection, developmental trajectories and extrinsic heterogeneity in Haldane’s rule. Evolution. 2015;69:2005–17.

21. Malone JH, Fontenot BE. Patterns of reproductive isolation in toads. PLOS ONE. 2008;3(12):e3900. doi: 10.1371/journal.pone.0003900.

22. Cutter AD, Bundus JD. Speciation and the developmental alarm clock. eLife. 2020;9:e56276. doi: 10.7554/eLife.56276.

23. Cutter AD. Speciation and development. Evol Dev. 2023;25:289–327.

24. Hernández-Hernández T, Miller EC, Román-Palacios C, Wiens JJ. Speciation across the Tree of Life. Biol Rev. 2021;96(4):1205–42. doi: 10.1111/brv.12698.

25. Wilson AC, Maxson LR, Sarich VM. Two types of molecular evolution. evidence from studies of interspecific hybridization. Proceedings of the National Academy of Sciences. 1974;71(7):2843–7. doi: 10.1073/pnas.71.7.2843.

26. Fitzpatrick BM. Rates of evolution of hybrid inviability in birds and mammals. Evolution. 2004;58(8):1865–70. doi: 10.1111/j.0014-3820.2004.tb00471.x.

27. Hedges SB, Marin J, Suleski M, Paymer M, Kumar S. Tree of Life reveals clock-like speciation and diversification. Mol Biol Evol. 2015;32(4):835–45. doi: 10.1093/molbev/msv037.

28. Corsi AK, Wightman B, Chalfie M. A transparent window into biology: a primer on *Caenorhabditis elegans*. Genetics. 2015;200(2):387–407. doi: 10.1534/genetics.115.176099.

29. Stevens L, Félix M-A, Beltran T, Braendle C, Caurcel C, Fausett S, et al. Comparative genomics of ten new Caenorhabditis species. Evolution Letters. 2019;3(2):217–36.

30. Vielle A, Callemeyn-Torre N, Gimond C, Poullet N, Gray JC, Cutter AD, et al. Convergent evolution of sperm gigantism and the developmental origins of sperm size variability in *Caenorhabditis* nematodes. Evolution. 2016;70(11):2485–503. doi: 10.1111/evo.13043. PubMed PMID: WOS:000387072900005.

31. Kiontke K, Felix M-A, Ailion M, Rockman M, Braendle C, Penigault J-B, et al. A phylogeny and molecular barcodes for *Caenorhabditis*, with numerous new species from rotting fruits. BMC Evol Biol. 2011;11(1):339. doi: 10.1186/1471-2148-11-339. PubMed PMID: doi:10.1186/1471-2148-11-339.

32. Kiontke K, Barriere A, Kolotuev I, Podbilewicz B, Sommer R, Fitch DH, et al. Trends, stasis, and drift in the evolution of nematode vulva development. Curr Biol. 2007;17(22):1925–37. PubMed PMID: 18024125.

33. Woodruff GC, Eke O, Baird SE, Felix MA, Haag ES. Insights into species divergence and the evolution of hermaphroditism from fertile interspecies hybrids of *Caenorhabditis* nematodes. Genetics. 2010;186:997–1012. Epub 2010/09/09. doi: genetics.110.120550 [pii] 10.1534/genetics.110.120550. PubMed PMID: 20823339.

34. Dey A, Jeon Y, Wang G-X, Cutter AD. Global population genetic structure of *Caenorhabditis remanei* reveals incipient speciation. Genetics. 2012;191(4):1257–69.

35. Li R, Ren X, Bi Y, Ho VWS, Hsieh C-L, Young A, et al. Specific down-regulation of spermatogenesis genes targeted by 22G RNAs in hybrid sterile males associated with an X-Chromosome introgression. Genome Res. 2016;26(9):1219–32. doi: 10.1101/gr.204479.116.

36. Felix MA, Braendle C, Cutter AD. A streamlined system for species diagnosis in *Caenorhabditis* (Nematoda: Rhabditidae) with name designations for 15 distinct biological species. PLoS ONE. 2014;9:e94723.

37. Cutter AD. X exceptionalism in *Caenorhabditis* speciation. Mol Ecol. 2018;27:3925–34. doi: 10.1111/mec.14423.

38. Baird SE, Seibert SR. Reproductive isolation in the Elegans-Group of *Caenorhabditis*. Nat Sci. 2013;5:18–25.

39. Baird SE, Yen W-C. Reproductive isolation in *Caenorhabditis*: terminal phenotypes of hybrid embryos. Evol Dev. 2000;2(1):9–15.

40. Baird SE, Sutherlin ME, Emmons SW. Reproductive isolation in Rhabditidae (Nematoda, Secernentea): mechanisms that isolate 6 species of 3 genera. Evolution. 1992;46(3):585–94. PubMed PMID: 1112.

41. Weadick CJ, Sommer RJ. Hybrid crosses and the genetic basis of interspecific divergence in lifespan in Pristionchus nematodes. J Evol Biol. 2017;30(3):650–7. doi: 10.1111/jeb.13022.

42. Kanzaki N, Ragsdale EJ, Herrmann M, Mayer WE, Sommer RJ. Description of three *Pristionchus* species (Nematoda: Diplogastridae) from Japan that form a cryptic species complex with the model organism *P. pacificus*. Zool Sci. 2012;29(6):403–17. doi: 10.2108/zsj.29.403.

43. Yoshida K, Herrmann M, Kanzaki N, Weiler C, Rödelsperger C, Sommer RJ. Two new species of *Pristionchus* (Nematoda: Diplogastridae) from Taiwan and the definition of the *pacificus* species-complex *sensu stricto*. J Nematol. 2018;50(3):355–68. Epub 2018/11/20. doi: 10.21307/jofnem-2018-019. PubMed PMID: 30451420; PubMed Central PMCID: PMCPMC6909367.

44. Xie D, Ma Y, Ye P, Liu Y, Ding Q, Huang G, et al. A newborn F-box gene blocks gene flow by selectively degrading phosphoglucomutase in species hybrids. Proceedings of the National Academy of Sciences. 2024;121(46):e2418037121. doi: 10.1073/pnas.2418037121.

45. Bloom J, Green R, Desai A, Oegema K, Rifkin SA. Hybrid incompatibility emerges at the one-cell stage in interspecies *Caenorhabditis* embryos. bioRxiv. 2024:2024.10.19.619171. doi: 10.1101/2024.10.19.619171.

46. Yoshida K, Rödelsperger C, Röseler W, Riebesell M, Sun S, Kikuchi T, et al. Chromosome fusions repatterned recombination rate and facilitated reproductive isolation during Pristionchus nematode speciation. Nat Ecol Evol. 2023;7(3):424–39. doi: 10.1038/s41559-022-01980-z.

47. Xie D, Ye P, Ma Y, Li Y, Liu X, Sarkies P, et al. Genetic exchange with an outcrossing sister species causes severe genome-wide dysregulation in a selfing *Caenorhabditis* nematode. Genome Res. 2022;32(11-12):2015–27. doi: 10.1101/gr.277205.122.

48. Yoshida K, Witte H, Hatashima R, Sun S, Kikuchi T, Röseler W, et al. Rapid chromosome evolution and acquisition of thermosensitive stochastic sex determination in nematode androdioecious hermaphrodites. Nat Comm. 2024;15(1):9649. doi: 10.1038/s41467-024-53854-6.

49. Sudhaus W, Kiontke K. Comparison of the cryptic nematode species *Caenorhabditis brenneri* sp. n. and *C. remanei* (Nematoda: Rhabditidae) with the stem species pattern of the *Caenorhabditis Elegans* group. Zootaxa. 2007;1456:45–62.

50. Turelli M, Moyle LC. Asymmetric postmating isolation: Darwin’s corollary to Haldane’s rule. Genetics. 2007;176(2):1059–88. Epub 2007/04/17. doi: genetics.106.065979 [pii] 10.1534/genetics.106.065979. PubMed PMID: 17435235; PubMed Central PMCID: PMC1894575.

51. Dey A, Jin Q, Chen Y-C, Cutter AD. Gonad morphogenesis defects drive hybrid male sterility in asymmetric hybrid breakdown of *Caenorhabditis* nematodes. Evol Dev. 2014;16:362–72.

52. Kozlowska JL, Ahmad AR, Jahesh E, Cutter AD. Genetic variation for post-zygotic reproductive isolation between *Caenorhabditis briggsae* and *Caenorhabditis* sp. 9. Evolution. 2012;66:1180–95.

53. Cutter AD. The polymorphic prelude to Bateson-Dobzhansky-Muller incompatibilities. Trends Ecol Evol. 2012;27:209–18.

54. Bi Y, Ren X, Yan C, Shao J, Xie D, Zhao Z. A genome-wide hybrid incompatibility landscape between *Caenorhabditis briggsae* and *C. nigoni*. PLoS Genet. 2015;11(2):e1004993. doi: 10.1371/journal.pgen.1004993.

55. Viswanath A, Cutter AD. Regulatory divergence as a mechanism for X-autosome incompatibilities in *Caenorhabditis* nematodes. Genome Biol Evol. 2023;15(4). doi: 10.1093/gbe/evad055. PubMed PMID: 37014784; PubMed Central PMCID: PMCPMC10147328.

56. Sánchez-Ramírez S, Weiss JG, Thomas CG, Cutter AD. Widespread misregulation of inter-species hybrid transcriptomes due to sex-specific and sex-chromosome regulatory evolution. PLoS Genet. 2021;17(3):e1009409. doi: 10.1371/journal.pgen.1009409.

57. Ting JJ, Woodruff GC, Leung G, Shin N-R, Cutter AD, Haag ES. Intense sperm-mediated sexual conflict promotes gametic isolation in *Caenorhabditis* nematodes. PLoS Biol. 2014;12:e1001915.

58. Ting JJ, Cutter AD. Demographic consequences of reproductive interference in multi-species communities. BMC Ecol. 2018;18:46.

59. Ting JJ, Tsai CN, Schalkowski R, Cutter AD. Genetic contributions to ectopic sperm cell migration in *Caenorhabditis* nematodes. G3: Genes|Genomes|Genetics. 2018;8(12):3891. doi: 10.1534/g3.118.200785.

60. Schalkowski R, Kasimatis KR, Greischar MA, Cutter AD. Reproductive interference alters species coexistence in nematodes due to asymmetric sperm-induced harm. Ecol Lett. 2025;28(1):e70067. doi: 10.1111/ele.70067.

61. Devi MP, Haryoso E, Rais EI, Karuniawan A, Yahya MQ, Richaud A, et al. Five new *Caenorhabditis* species from Indonesia provide exceptions to Haldane’s rule and partial fertility of interspecific hybrids. bioRxiv. 2025:2025.05.14.653126. doi: 10.1101/2025.05.14.653126.

62. Sloat SA, Noble LM, Paaby AB, Bernstein M, Chang A, Kaur T, et al. *Caenorhabditis* nematodes colonize ephemeral resource patches in neotropical forests. Ecology and Evolution. 2022;12(7):e9124. doi: 10.1002/ece3.9124.

63. Cutter AD. Divergence times in *Caenorhabditis* and *Drosophila* inferred from direct estimates of the neutral mutation rate. Mol Biol Evol. 2008;25(4):778–86. PubMed PMID: 18234705.

64. Dayi M, Kanzaki N, Sun S, Ide T, Tanaka R, Masuya H, et al. Additional description and genome analyses of *Caenorhabditis auriculariae* representing the basal lineage of genus *Caenorhabditis*. Scientific Reports. 2021;11(1):6720. doi: 10.1038/s41598-021-85967-z.

65. Sun S, Kanzaki N, Dayi M, Maeda Y, Yoshida A, Tanaka R, et al. The compact genome of *Caenorhabditis niphades* n. sp., isolated from a wood-boring weevil, *Niphades variegatus*. BMC Genomics. 2022;23(1):765. doi: 10.1186/s12864-022-09011-8.

66. Saxena AS, Salomon MP, Matsuba C, Yeh S-D, Baer CF. Evolution of the mutational process under relaxed selection in *Caenorhabditis elegans*. Mol Biol Evol. 2018;36(2):239–51. doi: 10.1093/molbev/msy213.

67. Charlesworth D. Don’t forget the ancestral polymorphisms. Heredity. 2010;105(6):509–10. doi: 10.1038/hdy.2010.14.

68. Li S, Jovelin R, Yoshiga T, Tanaka R, Cutter AD. Specialist versus generalist life histories and nucleotide diversity in Caenorhabditis nematodes. Proc R Soc B-Biol Sci. 2014;281(1777):20132858. doi: 10.1098/rspb.2013.2858.

69. Cutter AD, Morran LT, Phillips PC. Males, outcrossing, and sexual selection in *Caenorhabditis* nematodes. Genetics (Wormbook). 2019;213:27–57.

70. Kanzaki N, Tsai IJ, Tanaka R, Hunt VL, Liu D, Tsuyama K, et al. Biology and genome of a newly discovered sibling species of *Caenorhabditis elegans*. Nat Comm. 2018;9(1):3216. doi: 10.1038/s41467-018-05712-5.

71. dos Reis M, Thawornwattana Y, Angelis K, Telford Maximilian J, Donoghue Philip CJ, Yang Z. Uncertainty in the timing of origin of animals and the limits of precision in molecular timescales. Curr Biol. 2015;25(22):2939–50. doi: 10.1016/j.cub.2015.09.066.

72. Obbard DJ, Maclennan J, Kim K-W, Rambaut A, O’Grady PM, Jiggins FM. Estimating divergence dates and substitution rates in the *Drosophila* phylogeny. Mol Biol Evol. 2012;29(11):3459–73. doi: 10.1093/molbev/mss150.

73. Roux C, Fraïsse C, Romiguier J, Anciaux Y, Galtier N, Bierne N. Shedding light on the grey zone of speciation along a continuum of genomic divergence. PLoS Biol. 2016;14(12):e2000234. doi: 10.1371/journal.pbio.2000234.

74. Vásquez-Restrepo JD. A dataset containing speciation rates, uncertainty, and presence-absence matrices for more than 34,000 vertebrate species. bioRxiv. 2024:2024.04.09.588748. doi: 10.1101/2024.04.09.588748.

75. Dufresnes C, Brelsford A, Jeffries DL, Mazepa G, Suchan T, Canestrelli D, et al. Mass of genes rather than master genes underlie the genomic architecture of amphibian speciation. Proceedings of the National Academy of Sciences. 2021;118(36):e2103963118. doi: 10.1073/pnas.2103963118.

76. Dey A, Chan CK-W, Thomas CG, Cutter AD. Nucleotide hyperdiversity defines populations of *Caenorhabditis brenneri*. Proc Natl Acad Sci USA. 2013;110:11056–60.

77. Cutter AD, Jovelin R, Dey A. Molecular hyperdiversity and evolution in very large populations. Mol Ecol. 2013;22:2074–95.

78. Schiffman JS, Ralph PL. System drift and speciation. Evolution. 2021;76:236-51. doi: 10.1111/evo.14356.

79. Verster AJ, Ramani AK, McKay SJ, Fraser AG. Comparative RNAi screens in *C. elegans* and *C. briggsae* reveal the impact of developmental system drift on gene function. PLoS Genet. 2014;10(2):e1004077. doi: DOI 10.1371/journal.pgen.1004077. PubMed PMID: ISI:000332021500065.

80. Barrière A, Ruvinsky I. Pervasive divergence of transcriptional gene regulation in *Caenorhabditis* nematodes. PLoS Genet. 2014;10(6):e1004435. doi: 10.1371/journal.pgen.1004435.

81. Barkoulas M, Vargas Velazquez AM, Peluffo AE, Felix MA. Evolution of new *cis*-regulatory motifs required for cell-specific gene expression in *Caenorhabditis*. PLoS Genet. 2016;12(9):e1006278. doi: 10.1371/journal.pgen.1006278. PubMed PMID: 27588814; PubMed Central PMCID: PMCPMC5010242.

82. Valfort A-C, Launay C, Sémon M, Delattre M. Evolution of mitotic spindle behavior during the first asymmetric embryonic division of nematodes. PLoS Biol. 2018;16(1):e2005099. doi: 10.1371/journal.pbio.2005099.

83. Mallard F, Noble L, Guzella T, Afonso B, Baer CF, Teotónio H. Phenotypic stasis with genetic divergence. Peer Community Journal. 2023;3. doi: 10.24072/pcjournal.349.

84. Pybus OG, Harvey PH. Testing macro–evolutionary models using incomplete molecular phylogenies. Proc R Soc Lond B-Biol Sci. 2000;267(1459):2267–72. doi: 10.1098/rspb.2000.1278.

85. Peris D, Condamine FL. The angiosperm radiation played a dual role in the diversification of insects and insect pollinators. Nat Comm. 2024;15(1):552. doi: 10.1038/s41467-024-44784-4.

86. Benton MJ, Wilf P, Sauquet H. The Angiosperm Terrestrial Revolution and the origins of modern biodiversity. New Phytol. 2022;233(5):2017–35. doi: 10.1111/nph.17822.

87. Chaboureau A-C, Sepulchre P, Donnadieu Y, Franc A. Tectonic-driven climate change and the diversification of angiosperms. Proceedings of the National Academy of Sciences. 2014;111(39):14066–70. doi: 10.1073/pnas.1324002111.

88. Cutter AD. *Caenorhabditis* evolution in the wild. Bioessays. 2015;37(9):983–95. doi: 10.1002/bies.201500053.

89. Frézal L, Félix M-A. *C. elegans* outside the Petri dish. eLife. 2015;4:e05849. doi: 10.7554/eLife.05849. PubMed PMID: PMC4373675.

90. Rota-Stabelli O, Daley Allison C, Pisani D. Molecular timetrees reveal a Cambrian colonization of land and a new scenario for Ecdysozoan evolution. Curr Biol. 2013;23(5):392–8. doi: 10.1016/j.cub.2013.01.026.

91. Qing X, Zhang YM, Sun S, Ahmed M, Lo W-S, Bert W, et al. Phylogenomic insights into the evolution and origin of Nematoda. bioRxiv. 2023:2023.12.13.571554. doi: 10.1101/2023.12.13.571554.

92. Ahmed M, Roberts NG, Adediran F, Smythe AB, Kocot KM, Holovachov O. Phylogenomic analysis of the phylum Nematoda: Conflicts and congruences with morphology, 18S rRNA, and mitogenomes. Frontiers in Ecology and Evolution. 2022;9. doi: 10.3389/fevo.2021.769565.

93. Poinar GO. The Evolutionary History of Nematodes: Brill Academic Publishers; 2011.

94. Poinar Jr G, Kerp H, Hass H. *Palaeonema phyticum* gen. n., sp. n. (Nematoda: Palaeonematidae fam. n.), a Devonian nematode associated with early land plants. Nematology. 2008;10(1):9–14. doi: 10.1163/156854108783360159.

95. Laird CD, McConaughy BL, McCarthy BJ. Rate of fixation of nucleotide substitutions in evolution. Nature. 1969;224(5215):149–54. doi: 10.1038/224149a0.

96. Thomas JA, Welch JJ, Lanfear R, Bromham L. A generation time effect on the rate of molecular evolution in invertebrates. Mol Biol Evol. 2010;27(5):1173–80. doi: 10.1093/molbev/msq009.

97. Klass M, Hirsh D. Non-ageing developmental variant of *Caenorhabditis elegans*. Nature. 1976;260(5551):523–5.

98. Tellier A. Persistent seed banking as eco-evolutionary determinant of plant nucleotide diversity: novel population genetics insights. New Phytol. 2019;221(2):725–30. doi: 10.1111/nph.15424.

99. Nunney L. The effective size of annual plant populations: the interaction of a seed bank with fluctuating population size in maintaining genetic variation. Am Nat. 2002;160(2):195–204. doi: 10.1086/341017.

100. Vitalis R, Glémin S, Olivieri I. When genes go to sleep: the population genetic consequences of seed dormancy and monocarpic perenniality. Am Nat. 2004;163(2):295–311. doi: 10.1086/381041.

101. Lundemo S, Falahati-Anbaran M, StenØIen HK. Seed banks cause elevated generation times and effective population sizes of *Arabidopsis thaliana* in northern Europe. Mol Ecol. 2009;18(13):2798–811. doi: 10.1111/j.1365-294X.2009.04236.x.

102. Denver DR, Wilhelm LJ, Howe DK, Gafner K, Dolan PC, Baer CF. Variation in base-substitution mutation in experimental and natural lineages of *Caenorhabditis* nematodes. Genome Biol Evol. 2012;4(4):513–22. Epub 2012/03/23. doi: evs028 [pii] 10.1093/gbe/evs028. PubMed PMID: 22436997; PubMed Central PMCID: PMC3342874.

103. Konrad A, Brady MJ, Bergthorsson U, Katju V. Mutational landscape of spontaneous base substitutions and small indels in experimental *Caenorhabditis elegans* populations of differing size. Genetics. 2019;212(3):837–54. doi: 10.1534/genetics.119.302054.

104. Chen H-y, Krieg T, Mautz B, Jolly C, Scofield D, Maklakov AA, et al. Germline mutation rate is elevated in young and old parents in *Caenorhabditis remanei*. Evolution Letters. 2023;7(6):478–89. doi: 10.1093/evlett/qrad052.

105. Schulenburg H, Félix M-A. The natural biotic environment of *Caenorhabditis elegans*. Genetics. 2017;206(1):55.

106. Funk DJ, Nosil P, Etges WJ. Ecological divergence exhibits consistently positive associations with reproductive isolation across disparate taxa. P Natl Acad Sci USA. 2006;103(9):3209–13.

107. Stankowski S, Ravinet M. Defining the speciation continuum. Evolution. 2021;75(6):1256–73. doi: 10.1111/evo.14215.

108. Rosenblum EB, Sarver BAJ, Brown JW, Des Roches S, Hardwick KM, Hether TD, et al. Goldilocks meets Santa Rosalia: an ephemeral speciation model explains patterns of diversification across time scales. Evol Biol. 2012;39(2):255–61. doi: 10.1007/s11692-012-9171-x.

109. Cutter AD, Gray JC. Ephemeral ecological speciation and the latitudinal biodiversity gradient. Evolution. 2016;70(10):2171–85. doi: doi:10.1111/evo.13030.

110. Melander SL, Mueller RL. Comprehensive analysis of salamander hybridization suggests a consistent relationship between genetic distance and reproductive isolation across Tetrapods. Copeia. 2020;108(4):987–1003. doi: 10.1643/CH-19-319.

111. Gonzalez de la Rosa PM, Thomson M, Trivedi U, Tracey A, Tandonnet S, Blaxter M. A telomere-to-telomere assembly of *Oscheius tipulae* and the evolution of rhabditid nematode chromosomes. G3 Genes|Genomes|Genetics. 2021;11(1):jkaa020. doi: 10.1093/g3journal/jkaa020.

112. Stein LD, Bao Z, Blasiar D, Blumenthal T, Brent MR, Chen N, et al. The genome sequence of *Caenorhabditis briggsae*: A platform for comparative genomics. PLoS Biol. 2003;1(2):166–92. PubMed PMID: 14624247.

113. Orr HA, Turelli M. The evolution of postzygotic isolation: accumulating Dobzhansky-Muller incompatibilities. Evolution. 2001;55(6):1085–94. doi: 10.1111/j.0014-3820.2001.tb00628.x.

114. Thomas CG, Wang W, Jovelin R, Ghosh R, Lomasko T, Trinh Q, et al. Full-genome evolutionary histories of selfing, splitting and selection in *Caenorhabditis*. Genome Res. 2015;25:667–78.

115. Lamelza P, Ailion M. Cytoplasmic–nuclear incompatibility between wild isolates of *Caenorhabditis nouraguensis*. G3: Genes|Genomes|Genetics. 2017;7(3):823–34.

116. Ross JA, Koboldt DC, Staisch JE, Chamberlin HM, Gupta BP, Miller RD, et al. *Caenorhabditis briggsae* recombinant inbred line genotypes reveal inter-strain incompatibility and the evolution of recombination. PLoS Genet. 2011;7(7):e1002174. doi: 10.1371/journal.pgen.1002174. PubMed PMID: ISI:000293338600023.

117. Chung IW. A note on Haldane’s rule: hybrid inviability versus hybrid sterility. Evolution. 1992;46(5):1584–7. doi: 10.2307/2409965.

118. Wu CI, Davis AW. Evolution of postmating reproductive isolation: the composite nature of Haldane rule and its genetic bases. Am Nat. 1993;142(2):187–212. doi: Doi 10.1086/285534. PubMed PMID: ISI:A1993LU63300001.

119. Larson EL, Keeble S, Vanderpool D, Dean MD, Good JM. The composite regulatory basis of the large X-effect in mouse speciation. Mol Biol Evol. 2016;34(2):282–95. doi: 10.1093/molbev/msw243.

120. Cutter AD. Beyond Haldane’s rule: sex-biased hybrid dysfunction for all modes of sex determination. eLife. 2024;13:e96652.

121. Howe KL, Bolt BJ, Cain S, Chan J, Chen WJ, Davis P, et al. WormBase 2016: expanding to enable helminth genomic research. Nucleic Acids Res. 2016;44(D1):D774–D80. doi: 10.1093/nar/gkv1217.

122. Davis P, Zarowiecki M, Arnaboldi V, Becerra A, Cain S, Chan J, et al. WormBase in 2022—data, processes, and tools for analyzing Caenorhabditis elegans. Genetics. 2022;220(4):iyac003. doi: 10.1093/genetics/iyac003.

123. Sayers EW, Cavanaugh M, Clark K, Pruitt KD, Schoch CL, Sherry Stephen T, et al. GenBank. Nucleic Acids Res. 2022;50(D1):D161-D4. doi: 10.1093/nar/gkab1135.

124. Stevens L. Data associated with the Caenorhabditis Genomes Project (caenorhabditis.org). Zenodo; 2024. p. DOI: 10.5281/zenodo.12633738.

125. Fusca DD, Kasimatis KR, Zhu HV, Cutter AD. Dynamic birth and death of Argonaute gene family functional repertoire across *Caenorhabditis* nematodes. bioRxiv. 2024:2024.10. 27.620551.

126. Manni M, Berkeley MR, Seppey M, Simão FA, Zdobnov EM. BUSCO update: novel and streamlined workflows along with broader and deeper phylogenetic coverage for scoring of eukaryotic, prokaryotic, and viral genomes. Mol Biol Evol. 2021;38(10):4647–54. doi: 10.1093/molbev/msab199.

127. Loytynoja A, Goldman N. Phylogeny-aware gap placement prevents errors in sequence alignment and evolutionary analysis. science. 2008;320(5883):1632–5.

128. Minh BQ, Schmidt HA, Chernomor O, Schrempf D, Woodhams MD, von Haeseler A, et al. IQ-TREE 2: new models and efficient methods for phylogenetic inference in the genomic era. Mol Biol Evol. 2020;37(5):1530–4. doi: 10.1093/molbev/msaa015.

129. Kalyaanamoorthy S, Minh BQ, Wong TK, Von Haeseler A, Jermiin LS. ModelFinder: fast model selection for accurate phylogenetic estimates. Nature methods. 2017;14(6):587–9.

130. Zhang C, Mirarab S. ASTRAL-Pro 2: ultrafast species tree reconstruction from multi-copy gene family trees. Bioinformatics. 2022;38(21):4949–50.

131. Emms DM, Kelly S. OrthoFinder: phylogenetic orthology inference for comparative genomics. Genome Biol. 2019;20(1):238. doi: 10.1186/s13059-019-1832-y.

132. Sela I, Ashkenazy H, Katoh K, Pupko T. GUIDANCE2: accurate detection of unreliable alignment regions accounting for the uncertainty of multiple parameters. Nucleic Acids Res. 2015;43(W1):W7–W14.

133. Tamura K, Stecher G, Kumar S. MEGA11: molecular evolutionary genetics analysis version 11. Mol Biol Evol. 2021;38(7):3022–7.

134. Bromham L, Penny D. The modern molecular clock. Nat Rev Genet. 2003;4(3):216–24. doi: 10.1038/nrg1020.

135. Cutter AD, Wasmuth J, Blaxter ML. The evolution of biased codon and amino acid usage in nematode genomes. Mol Biol Evol. 2006;23:2303–15. PubMed PMID: 16936139.

136. Duret L, Mouchiroud D. Expression pattern and, surprisingly, gene length shape codon usage in Caenorhabditis, Drosophila, Arabidopsis. Proc Natl Acad Sci USA. 1999;96(8):4482–7. PubMed PMID: 10200288.

137. Kosakovsky Pond SL, Poon AF, Velazquez R, Weaver S, Hepler NL, Murrell B, et al. HyPhy 2.5—a customizable platform for evolutionary hypothesis testing using phylogenies. Mol Biol Evol. 2020;37(1):295–9.

138. Phillips MJ. Branch-length estimation bias misleads molecular dating for a vertebrate mitochondrial phylogeny. Gene. 2009;441(1):132–40. doi: 10.1016/j.gene.2008.08.017.

139. Bouckaert R, Vaughan TG, Barido-Sottani J, Duchêne S, Fourment M, Gavryushkina A, et al. BEAST 2.5: An advanced software platform for Bayesian evolutionary analysis. PLoS Comp Biol. 2019;15(4):e1006650.

140. Douglas J, Jiménez-Silva CL, Bouckaert R. StarBeast3: adaptive parallelized Bayesian inference under the multispecies coalescent. Syst Biol. 2022;71(4):901–16.

141. R Core Team. R: A Language and Environment for Statistical Computing. Vienna, Austria: R Foundation for Statistical Computing; 2024.

142. Rambaut A, Drummond AJ, Xie D, Baele G, Suchard MA. Posterior summarization in Bayesian phylogenetics using Tracer 1.7. Syst Biol. 2018;67(5):901–4.

143. Jonasson J, Harkonen T, Sundqvist L, Edwards SV, Harding KC. A unifying framework for estimating generation time in age-structured populations: implications for phylogenetics and conservation biology. Am Nat. 2022;200(1):48–62. doi: 10.1086/719667.

144. Bapst DW. paleotree: an R package for paleontological and phylogenetic analyses of evolution. Meth Ecol Evol. 2012;3(5):803–7.

145. Revell LJ. phytools 2.0: an updated R ecosystem for phylogenetic comparative methods (and other things). PeerJ. 2024;12:e16505. doi: 10.7717/peerj.16505.

146. Smith ML, Hahn MW. New approaches for inferring phylogenies in the presence of paralogs. Trends Genet. 2021;37(2):174–87. doi: 10.1016/j.tig.2020.08.012.

147. Smith ML, Vanderpool D, Hahn MW. Using all gene families vastly expands data available for phylogenomic inference. Mol Biol Evol. 2022;39(6):msac112. doi: 10.1093/molbev/msac112.

148. Smith ML, Hahn MW. Using paralogs for phylogenetic inference mitigates the effects of long-branch attraction. bioRxiv. 2024:2024.12.06.627281. doi: 10.1101/2024.12.06.627281.

149. Dolgin ES, Charlesworth B, Baird SE, Cutter AD. Inbreeding and outbreeding depression in *Caenorhabditis* nematodes. Evolution. 2007;61:1339–52. PubMed PMID: 17542844.

150. Sobel JM, Chen GF. Unification of methods for estimating the strength of reproductive isolation. Evolution. 2014;68(5):1511–22. doi: 10.1111/evo.12362.

151. Alfieri JM, Hingoranee R, Athrey GN, Blackmon H. Domestication is associated with increased interspecific hybrid compatibility in landfowl (order: Galliformes). J Hered. 2024;115(1):1–10. doi: 10.1093/jhered/esad059.

